# Seasonal plasticity in morphology and metabolism differs between migratory North American and resident Costa Rican monarch butterflies

**DOI:** 10.1101/2022.06.17.495480

**Authors:** Ayşe Tenger-Trolander, Cole R. Julick, Wei Lu, Delbert André Green, Kristi L. Montooth, Marcus R. Kronforst

## Abstract

Environmental heterogeneity in temperate latitudes is expected to maintain seasonally plastic life-history strategies that include the tuning of morphologies and metabolism that support overwintering. For species that have expanded their ranges into tropical latitudes, it is unclear the extent to which the capacity for plasticity will be maintained or will erode with disuse. The migratory generations of the North American (NA) monarch butterfly *Danaus plexippus* lead distinctly different lives from their summer generation NA parents and their tropical descendants living in Costa Rica (CR). NA migratory monarchs postpone reproduction, travel thousands of kilometers south to overwinter in Mexico, and subsist on little food for months. Whether recently dispersed populations of monarchs such as those in Costa Rica, which are no longer subject to selection imposed by migration, retain ancestral seasonal plasticity is unclear. To investigate differences in seasonal plasticity, we reared NA and CR monarchs in summer and autumn in Illinois, USA, and measured seasonal reaction norms for aspects of morphology and metabolism related to flight. NA monarchs were seasonally plastic in forewing and thorax size, increasing wing area and thorax to body mass ratio in autumn. While CR monarchs increased thorax mass in autumn, they did not increase the area of the forewing. NA monarchs maintained similar resting and maximal flight metabolic rates across seasons. However, CR monarchs had elevated metabolic rates in autumn. Our findings suggest that the recent expansion of monarchs into habitats that support year-round breeding may be accompanied by (1) the loss of some aspects of morphological plasticity as well as (2) the underlying physiological mechanisms that maintain metabolic homeostasis in the face of temperature heterogeneity.

## Introduction

Fluctuating seasonal environments in temperate habitats are expected to favor the evolution of overwintering strategies that are, by their nature, plastic responses to the environment (Moran 1992; Kingsolver and Huey 1998). For some species, these strategies involve changes in physiology and morphology that accompany overwintering in place, while for other species they involve physiological and morphological changes that support seasonal migration (Arnold et al. 2004; Butler and Woakes 2001). Theory for the evolutionary maintenance and loss of plasticity has been well developed (Via and Lande 1985, 1987; de Jong 1990; Van Tienderen 1991; Gomulkiewicz and Kirkpatrick 1992; Moran 1992; Gavrilets and Scheiner 1993) and there are established empirical frameworks and organismal systems for investigating the evolutionary dynamics of seasonal plasticity (Kingsolver and Huey 1998; Scheiner 1993). Trait plasticity may be lost when species ranges expand out of temperate, seasonal environments and into tropical, constant environments. This loss may occur via costs of plasticity that include a reduced efficacy of selection (Van Tienderen 1991; Kawecki 1994; Whitlock 1996; DeWitt et al. 1998; Van Dyken and Wade 2010) or genetic assimilation whereby adaptation to a new constant environment fixes the trait value (Waddington 1961; Price et al. 2003; Lande 2009; Schleicherová et al. 2013; Wan et al. 2018).

Trait plasticity may be a key factor enabling population persistence as environments across the globe become more variable (Catullo et al. 2019; Matesanz and Ramirez-Valiente 2019; O’Connor et al. 2012; Price et al. 2003; Sgro et al. 2016), motivating a better empirical understanding of how and when plasticity is lost. The loss of plasticity through assimilation, where some populations evolve a fixed phenotype that maximizes fitness in a new, stable environment (Aubret and Shine 2009; Corl et al. 2018) has been demonstrated in a handful of systems, including at a mechanistic and genetic level for wing coloration in common buckeye butterflies (van der Burg et al. 2020). A series of studies by Cooper et al. (2012, 2014) suggests that cellular membrane plasticity in phospholipid composition can be eroded through disuse or via costs of maintaining plasticity in fruit fly populations evolved in constant thermal environments. Relaxed selection at cold-acclimation genes coincides with range expansion into warmer latitudes in *Arabidopsis* (Zhen and Ungerer 2008; Zhen et al. 2011). However, seasonal plasticity is complex in that it involves suites of traits to support divergent physiologies or life histories across seasons (Williams et al. 2017; Wilsterman 2021).Investigating how this complex multi-trait plasticity is lost (e.g., piecemeal versus wholesale loss) when species ranges expand into less seasonal latitudes may provide insight into both the mechanisms of trait integration and trait loss, as well as identify aspects of seasonal plasticity that may be retained and respond to different environmental cues in the new environment.

North American (NA) monarch butterflies (*Danaus plexippus*) are well known for their long-distance seasonal migration plasticity that meets different dispersal, reproductive, and energetic demands across generations (Reppert and de Roode 2018). Summer generations are short-lived and reproduce shortly after adult eclosion. The autumn/winter generation lives for 8-12 months during which they migrate to their overwintering grounds in Mexico where they remain in reproductive diapause until the following spring. They then migrate back into the Southern United States and successive generations recolonize northern latitudes. In N. America, seasonally variable environmental conditions are predicted to maintain plasticity for many aspects of morphology and physiology that support the different life-history strategies in summer versus autumn/winter generations. Notably, previous studies comparing summer (non-migratory) and autumn (migratory) generation monarchs have shown that NA monarchs eclose in reproductive diapause, have increased longevity and cold tolerance, greater fat stores, differences in sun compass neuropil volume, and a strong drive to fly south in autumn compared with summer-eclosing monarchs (Barker and Herman 1976; Herman and Tatar 2001; Goehring and Oberhauser 2002; Brower et al. 2006; Zhu et al. 2008, 2009; Heinze et al. 2013; Tenger-Trolander et al. 2019).

NA monarchs have expanded their range through multiple independent dispersal events into tropical latitudes that lack seasonal heterogeneity and support resident, year-round breeding populations (Zhan et al. 2014), making monarchs a good system to investigate the loss of complex multi-trait plasticity. These populations are descendants of the migratory NA population, but have lost long-distance migratory behavior and differ in two migration-relevant phenotypes – wing size and sun compass neuron tuning to sunlight (Altizer and Davis 2010; Freedman et al. 2020; Nguyen et al. 2021). Today, non-migratory populations can be found in Central and South America, the Caribbean, the Iberian Peninsula, Morocco, the Pacific Islands, Australia, and New Zealand (Zhan et al. 2014; Pfeiler et al. 2017). In Australia, there are both migratory and non-migratory populations that exhibit plasticity in reproductive development (James 1984; Dingle et al. 1999; Freedman et al. 2018), suggesting that some aspects of seasonal migration plasticity may be maintained within some dispersed populations.

Here, we quantified plasticity across generations in NA monarchs for wing morphology and metabolic traits that are related to long-distance migration, and then asked whether Costa Rican (CR) monarch butterflies have lost or decreased plasticity in these traits (Figure 1). Using a common garden experiment with seasonal rearing of NA and CR monarchs, we tested the prediction that migratory populations have greater plasticity in response to seasonal rearing conditions than do non-migratory populations that no longer experience temperate-latitude seasonality in temperature, day length, and host-plant availability (Figure 1).

**Figure 1.**
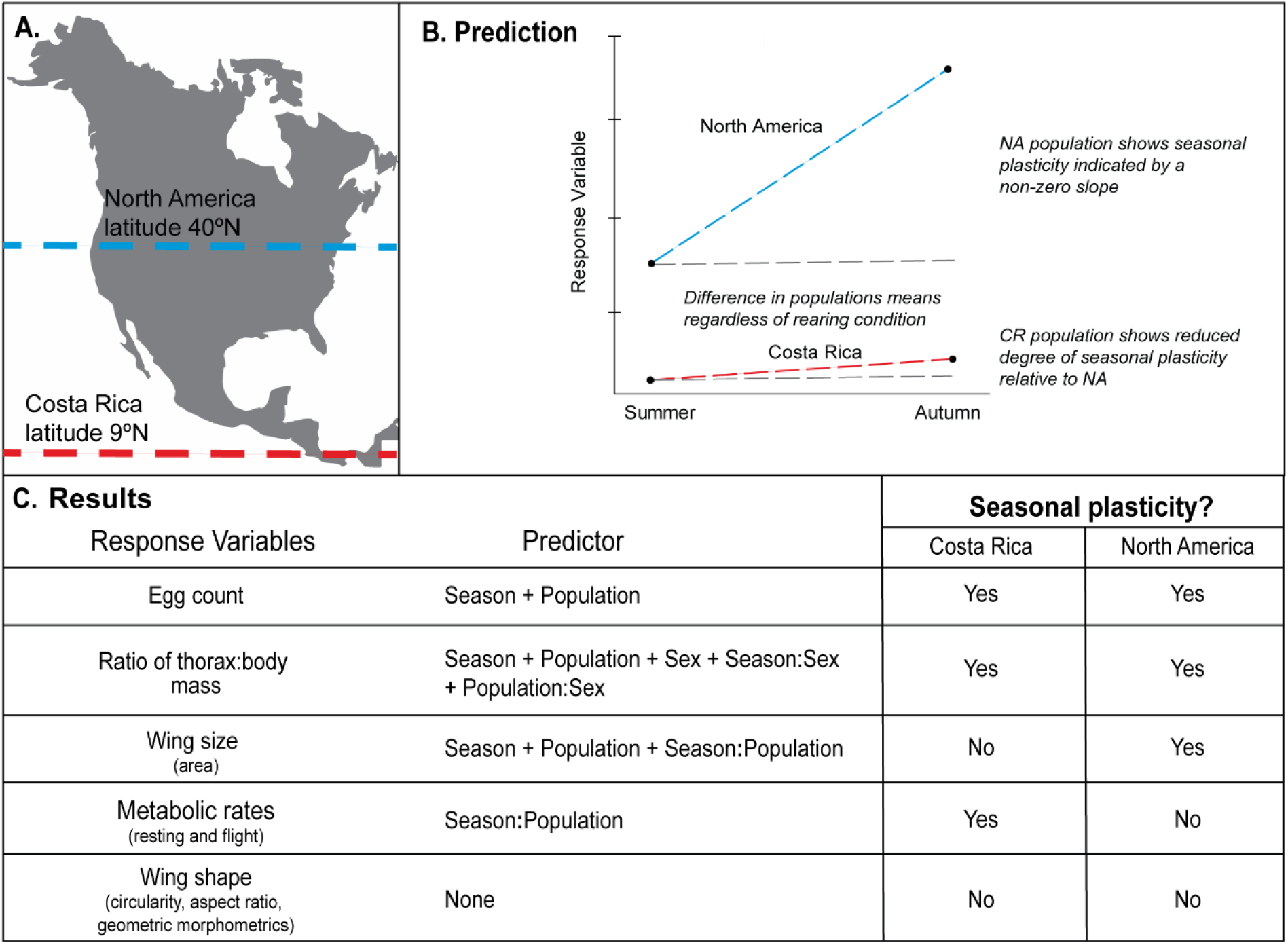
Summary of the experiment, predictions, and main findings investigating seasonal plasticity in Costa Rican (CR) and North American (NA) monarchs. A) Map of North and Central America indicating NA (blue) and CR (red) monarch respective latitudes. B) Prediction and potential outcomes of a possible response variable (i.e., a trait). Populations may differ in trait value regardless of seasonal rearing condition. Populations may also differ in seasonal trait plasticity. A non-zero reaction norm between summer and autumn trait values within a population indicates the presence of seasonal plasticity (blue and red vs grey lines), and differences in reaction norm between populations indicate differences in degree of plasticity (different slope of blue and red line). C) List of traits measured in this study, the independent variables (population, rearing season, and sex) that explained significant variance in the trait, and whether each population exhibited seasonal trait plasticity.

## Methods

### Trait Selection

We selected physiological and morphological traits that are likely subject to different selective pressures depending upon seasonal environmental heterogeneity, including egg count, thorax and abdomen mass, wing size, wing shape, and metabolic rate. Plasticity in these traits is thought to impact monarch success in reproduction, migration and overwintering. We counted the number of mature oocytes as a measure of reproductive arrest, a well-known phenomenon in migrating NA monarchs that is correlated with their longevity and overwintering strategy (Barker and Herman 1976; Goehring and Oberhauser 2002). We measured resting and maximal flight metabolic rates to quantify plasticity in energy demand that supports adult maintenance and flight. We massed the thorax and abdomen separately to estimate mass associated with the flight muscles and with the reproductive organs and fat body respectively. We measured forewing size and shape as traits associated with flight efficiency. Larger wings generate more lift due to lower wing loading, and more narrow wings with high aspect ratios decrease drag. (Winkler and Leisler 1992; Senar et al. 1994; Lockwood et al. 1998; Swaddle and Witter 1998; Dudley 2000; Egbert and Belthoff 2003; Wang 2004).

### Seasonal rearing

We reared two generations (summer and autumn) of NA and CR monarch butterflies outdoors in Chicago, IL in 2016 and 2017. Rearing was done under permits from USDA-APHIS. Butterflies that emerged in July and August were designated as the summer generation and those that emerged in September and October were designated as the autumn generation. In both years, the autumn generation were the offspring of the summer generation. In 2016, we measured metabolic traits in female and male adult monarchs and counted the number of mature oocytes in the females. We repeated these measurements for monarchs reared in 2017, with the addition of morphological measurements, including body mass, forewing size, and forewing shape.

### Sample sizes

We reared 576 monarchs (149 individuals in 2016 and 427 individuals in 2017) and measured 573 of these for at least one trait. We measured 179 individual’s metabolic rates, dissected 165 females to count the number of mature oocytes present in the abdomen, dried and massed 184 individuals, assayed geometric morphometric shape traits of 254 individuals, and measured the forewing size and shape traits for 237 individuals. Some individuals were used for multiple measurements, but those for which we counted oocytes could not be used for mass measurements and vice versa. In addition, butterflies for which we measured metabolic traits were more likely to be tattered and excluded from wing trait measurements. Further details on the number of individuals measured for each trait by rearing year, season of development, and population can be found in Table 1.

**Table 1.**
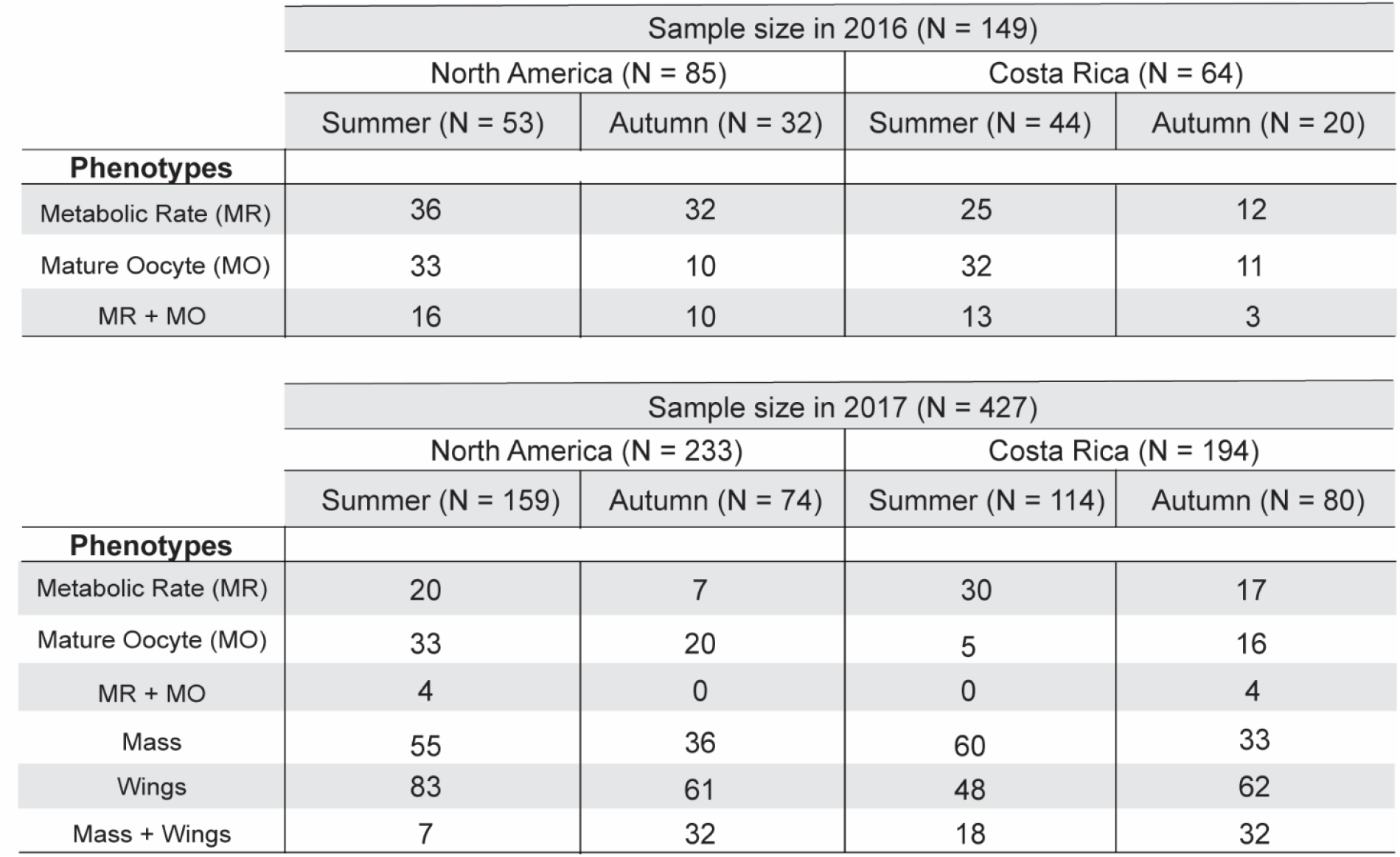
Sample sizes for all traits measured. Top) For samples reared in 2016, we assayed metabolic rate (MR) and number of mature oocytes (MO). MR + MO indicates the subset of individuals from the MR and MO row tallies for which we have measurements of both traits for the same individual. Bottom) For samples reared in 2017, in addition to metabolic rate assays and oocyte counts, we measured morphological traits using wings and bodies. MR + MO indicates the subset of individuals from the MR and MO row tallies for which we have measurements of both traits for the same individual, and Mass + Wings indicates the subset of individuals from the Mass and Wings row tallies for which we have measurements of both traits for the same individual.

|  | Sample size in 2016 (N = 149) |  |  |  |
| --- | --- | --- | --- | --- |
|  | North America (N = 85) |  | Costa Rica (N = 64) |  |
|  | Summer (N = 53) | Autumn (N = 32) | Summer (N = 44) | Autumn (N = 20) |
| Phenotypes |  |  |  |  |
| Metabolic Rate (MR) | 36 | 32 | 25 | 12 |
| Mature Oocyte (MO) | 33 | 10 | 32 | 11 |
| MR + MO | 16 | 10 | 13 | 3 |

|  | Sample size in 2017 (N = 427) |  |  |  |
| --- | --- | --- | --- | --- |
|  | North America (N = 233) |  | Costa Rica (N = 194) |  |
|  | Summer (N = 159) | Autumn (N = 74) | Summer (N = 114) | Autumn (N = 80) |
| Phenotypes |  |  |  |  |
| Metabolic Rate (MR) | 20 | 7 | 30 | 17 |
| Mature Oocyte (MO) | 33 | 20 | 5 | 16 |
| MR + MO | 4 | 0 | 0 | 4 |
| Mass | 55 | 36 | 60 | 33 |
| Wings | 83 | 61 | 48 | 62 |
| Mass + Wings | 7 | 32 | 18 | 32 |

### Genetic composition

We derived CR monarchs from ∼20 pupae obtained from a butterfly breeder in Costa Rica in 2016 and in 2017. While Central and South American monarchs remain the most genetically similar to NA monarchs, perhaps as a result of continuing gene flow (Pierce et al. 2014; Freedman et al. 2020), their estimated divergence time of 2,000-3,000 years from the N. American population is the largest among the three dispersals (Zhan et al. 2014).

All NA monarchs reared in 2017 were derived from wild-caught NA monarchs captured in Chicago, IL and morphological traits were only measured in 2017 (Table 1). The NA monarchs reared in 2016 were derived from two sources: wild-caught and commercially sourced NA individuals. At the time of the first common garden experiment, we were not aware of the genetic distinctiveness of the commercial NA lineage compared with the wild NA population (Tenger-Trolander et al. 2019). Of the 179 individuals assayed for MR (Supplemental Information, Table S1), only 18 individuals were purely commercial (15 summer-reared and 3 autumn-reared), 21 were NA/Commercial F1 crosses reared in summer, and 29 were backcrosses (NA/Commercial F1 backcrossed to NA) reared in autumn. While a proportion of pure commercial NA monarchs have lost the propensity to orient south in response to autumn rearing conditions (Tenger-Trolander et al. 2019; Tenger-Trolander and Kronforst 2020), the backcrosses (NA/Commercial F1 backcrossed to NA) reared in autumn showed clear southern orientation (Tenger-Trolander et al. 2019). The commercial lineage also enters reproductive diapause when reared outdoors in autumn implying some physiological responses in the wild and commercial NA populations are similar (Tenger-Trolander et al. 2019).

### Animal Husbandry

We housed the monarchs from their respective populations in medium size (91.5cm x 30.5cm^2^) mesh pop-up cages outdoors with access to the host plant, *Asclepias syriaca*. After females laid eggs, we transferred the eggs to small (30.5cm^3^) outdoor mesh pop-up cages and fed larvae on a diet of wild-collected *A. syriaca* cuttings. All pop-up cages were contained inside two large outdoor 1.83m^3^ mesh cages separated by population of origin. As individuals eclosed, they were collected as virgin males and females, labeled with a unique ID, and left outdoors for a minimum of three days. Adults were then either shipped to Lincoln, NE for metabolic measurement or measured for morphological traits in Chicago. Adult butterflies were shipped overnight in glassine envelopes, spending between 12-24 hours in a dark cardboard box. Upon arrival, butterflies were separated by sex and housed in large collapsible butterfly cages in a laboratory space with natural light. Prior to metabolic measurements, all individuals were given at least 48 hours to acclimate. Individuals kept in Chicago were also separated by sex and housed outdoors in Chicago, IL until frozen for morphological trait measurements. All butterflies had access to a constant supply of artificial nectar (Birds Choice Butterfly Nectar, Chilton, WI).

In 2017, summer-reared monarchs were shipped back to Chicago from Lincoln to found the autumn generation due to a summer die-off caused by the spillover of the pesticide permethrin from a neighboring yard in Chicago. None of the summer-reared monarchs that we measured were present when this exposure occurred, and the subsequent autumn generation of monarchs was founded by individuals not present during the die-off.

### Mature oocyte counts

Females were kept separately from males and never mated. We dissected females by making a longitudinal cut down the abdomen to remove eggs. We then counted the number of mature oocytes present. Immature and mature oocytes in monarchs are distinguished by the shape of the chorion. A smooth chorion surface indicates an immature oocyte while a chorion with ridges is considered mature.

### Body mass measurements

We removed the wings, antennae, head, and legs from the body. We separated the thorax and abdomen and dried them at 60°C in an incubator with a silica crystal desiccant for 72 hours. After drying, we weighed the thorax and abdomen both separately and together on an analytical balance.

### Wing size and shape measurements

We placed a single forewing and hindwing on a sheet of gridded paper with 0.635 cm squares or on a white sheet of paper with a metric ruler in view. We photographed the wings using a DSLR Canon EOS 70d camera with an 18-55mm lens. We scaled each photo by number of pixels/cm and converted the color photos to 8-bit black/white images in ImageJ (Schindelin et al. 2012; Rueden et al. 2017). We filled in non-black portions of the forewing with black to measure area (Supplementary Information, Figure S1). Before measuring area or shape attributes, we smoothed the contours of the forewings with the ImageJ plugin ‘Shape smoothing’ (Erdenetsogt and Wagner 2016). ‘Shape smoothing’ applies a Fourier transformation to gain Fourier descriptors (FDs). We kept 0.35% of FDs relative to the total number of FDs identified in the image (Supplementary Information, Figure S1). We then measured the area (in cm^2^), aspect ratio (length/width), and circularity (4π*area/perimeter^2^) of each forewing in ImageJ (Rueden et al. 2017). To measure aspect ratio, ImageJ finds the longest length (major axis) and width (minor axis) of the object while maintaining the perpendicular intersection of both lines and divides the major axis length by the minor. Higher circularity scores indicate a more circular wing shape whereas lower scores more polygonal or angular shapes. Circularity is different than roundness (4*area/(π*major_axis^2^)). For example, a hexagon has high circularity and low roundness whereas an oval has low circularity and high roundness.

We also used 2D landmark-based geometric morphometrics to assess shape differences. Using the software tpsDIG2ws, we placed 16 landmarks at homologous points (vein intersections and margins) on each forewing (Rohlf 2006) (Supplementary Information, Figure S2). We analyzed the resulting landmark data in R using the package ‘Geomorph’ (Adams and Collyer 2020). We performed a general Procrustes analysis that removed differences in orientation and size, allowing us to focus exclusively on shape differences (Supplementary Information, Figure S3). We then calculated the mean shape which is the average landmark coordinates for a set of aligned wings. For each of the 16 landmarks, we calculated the distance between the individual’s coordinates and the group mean coordinates and summed those distances to find each specimen’s total distance from the mean shape.

### Metabolic rate measures

Using flow-through respirometry, we estimated resting or routine metabolic rate (MR) and maximal flight MR from the volume of CO_2_ (VCO_2_) produced by individual adult monarchs ranging from 3-45 days old. While not ideal, this age range was a consequence of shipping logistics between Chicago and Lincoln. Age was not a significant predictor of MR, even when controlling for mass in an analysis of covariance (routine MR, *P* = 0.157; flight MR, *P*=0.310). Older butterflies tended to be smaller (effect of age on mass, *P* = 0.009), and variation in mass was accounted for in our statistical analysis of MR (see below in *Statistical Analyses)*. Butterflies were placed in a 3.3-liter glass cylindrical container covered with a piece of black velvet cloth ensuring complete darkness during resting MR measurements. CO_2_-free, dry air was pumped through the container at a rate of 3 liters/minute using a dual pump system (Sable Systems International, Las Vegas, NV, USA) coupled with a mass-flow valve (Sierra Instruments, Monterey, CA, USA). After the air left the measurement container, it was subsampled at 100 ml/min using a SS-4 Sub-Sampler pump (Sable Systems International, Las Vegas, NV, USA), scrubbed of water and then passed into a high-performance CO_2_/H_2_O differential gas analyzer (LI-7000, Li-Cor, Lincoln, NE, USA) to quantify CO_2_. All MR data were collected using the Expedata software package (Sable Systems International, Las Vegas, NV, USA).

Individuals rested in the cloth-covered chamber at 21°C for a minimum of 25 minutes prior to metabolic rate measurement. Before removing the cloth, we recorded resting MR until a stable resting MR was established. We then removed the cloth and exposed the individual to full-spectrum UV light. After 30 seconds of light exposure, we induced flight by gently shaking the container. We recorded 10 minutes of CO_2_ production during flight. If butterflies stopped flying during this 10-min period, we gently shook the chamber to induce flight. After flying for 10 minutes, we turned off the light and covered the container to allow the butterfly to return to a stable resting MR. While 21°C is cooler than others’ have used to measure flight metabolic rate (e.g., Zhan et al. 2014; Pocius et al. 2022), we chose this because it was the common garden temperature at which all monarchs were being held in the lab prior to measurements. We experienced no issues inducing flight, which was likely facilitated by warming due to the full-spectrum UV light and monarch thermoregulatory behavior (Masters et al. 1988). The maximal rate of CO_2_ production sustained over a 1-min period during the 10-minute flight was used as our estimate of maximal flight MR. Before and after each metabolic measurement, baseline CO_2_ values were recorded and drift-corrected using the two-endpoint method in Expedata. To compensate for a response lag in the respirometry system, we utilized the “Z-transformation” function (instantaneous transformation) in Expedata. Raw CO_2_ values were converted from parts per million to ml/hr.

### Statistical analyses

For analyses of morphological traits, we used the R package ‘glmulti’ to automatically select the best fit generalized linear model for each trait and determine which of the independent variables (sex, population, season) were significant predictors of the measurements (Calcagno and de Mazancourt, 2010; R Core Team 2013). We fit thorax mass (grams), thorax:body mass ratio, forewing area (cm^2^), and abdomen mass (log-transformed) within the Gaussian family as these traits were normally distributed (Supplementary Information, Figure S4A, Table S2 and S3). For egg counts, we fit the model with a negative binomial distribution (Supplementary Information, Figure S4B and Table S4).

To quantify the association between each independent variable and dependent variable for our glms, we calculated effect size with the statistic Eta squared (η2) which is the ratio of each group’s sum of squares to the total sum of squares. It is interpreted as the percentage of variance accounted for by each variable in the glm. For the negative binomial model, we relied on model coefficients to determine the predominant effect. We further performed the rank-based nonparametric Kruskal-Wallis test to determine whether there were differences between groups and then a post hoc Dunn test (with a Bonferroni correction for multiple testing) to determine which groups were different. Circularity scores, aspect ratios, and mean shape distances were not normally distributed, and various transformations of the measurements did not yield normal distributions (Supplementary Information, Figure S4C). In these cases, we relied on the rank-based nonparametric Kruskal-Wallis test with post hoc Dunn test (with a Bonferroni correction) to identify differences between groups.

We used standardized major axis regression (SMA) implemented in the R package “smatr” (Warton et al. 2006; R Core Team 2013) to test for the effects of rearing conditions and population on metabolic rate. SMA controls for the relationship between metabolic rate and mass (in our case, whole-body wet mass) like an analysis of covariance, but it accounts for the fact that both metabolic rate and mass are measured with error. We used SMA to fit the metabolic scaling relationship between ln(VCO_2_) and ln(mass) for all individuals within particular combinations of the factors sex, population (NA and CR), and rearing season (summer and autumn). We first tested whether the scaling relationship between MR and mass was similar between levels of our independent factors (i.e., testing for difference in slopes). When there was no evidence of a mass x factor interaction, we fit a common slope and then tested whether there was a significant effect of the factor on the elevation of the relationship between MR and mass (i.e., a difference in the mass-specific metabolic rate) or a significant shift along the x-axis between factor levels (i.e., a difference in mass). Because females and males did not differ significantly in the scaling relationship between MR and mass or in mass-specific MR (Supplementary Information, Table S5), the sexes were combined for all subsequent analyses. The scaling exponents relating ln(VCO_2_) and ln(mass) were greater than 1. While this deviates from the broad interspecific pattern where metabolic rate scales with mass to the ¾ power, intraspecific scaling exponents frequently deviate from this expectation (Glazier 2005; Greenlee et al. 2014). These scaling exponents may also have differed if we had chosen a different measure for mass, e.g., with wings removed.

We used mass-corrected resting and flight MRs to test for the statistical significance of interactions between population and rearing condition on metabolic rates, which would indicate a difference in plasticity between NA and CR monarchs. Mass-corrected MRs were obtained by taking the residual for each individual from the SMA fits of ln(VCO_2_) as a function of ln(mass) and adding back the average ln(VCO_2_) to obtain a meaningful scale as in Hoekstra et al. (2013). The mass-corrected MRs were used as the dependent variable in linear models to test for the effects of population, rearing season, and the statistical interaction between population and rearing season.

## Results

### Both NA and CR monarchs exhibit seasonal plasticity in female reproduction

The number of mature oocytes present in a female monarch’s abdomen was best explained by the additive effects of season and population (Supplementary Information, Table S4). Both CR and NA monarchs had fewer mature oocytes when reared in autumn (Figure 2, Kruskal-Wallis χ^2^ = 64.32, df = 3, *P* = 7.02e-14), consistent with the known seasonal reproductive diapause. Season had a larger effect size estimate (1.31, 95% CI [0.94, 1.66]) than did population (−0.62, 95% CI [-0.97, -0.28]) with little CI overlap. Although our model suggests there was a significant effect of population, differences between CR and NA monarchs within each seasonal rearing environment were not significant after correcting for multiple tests.

**Figure 2.**
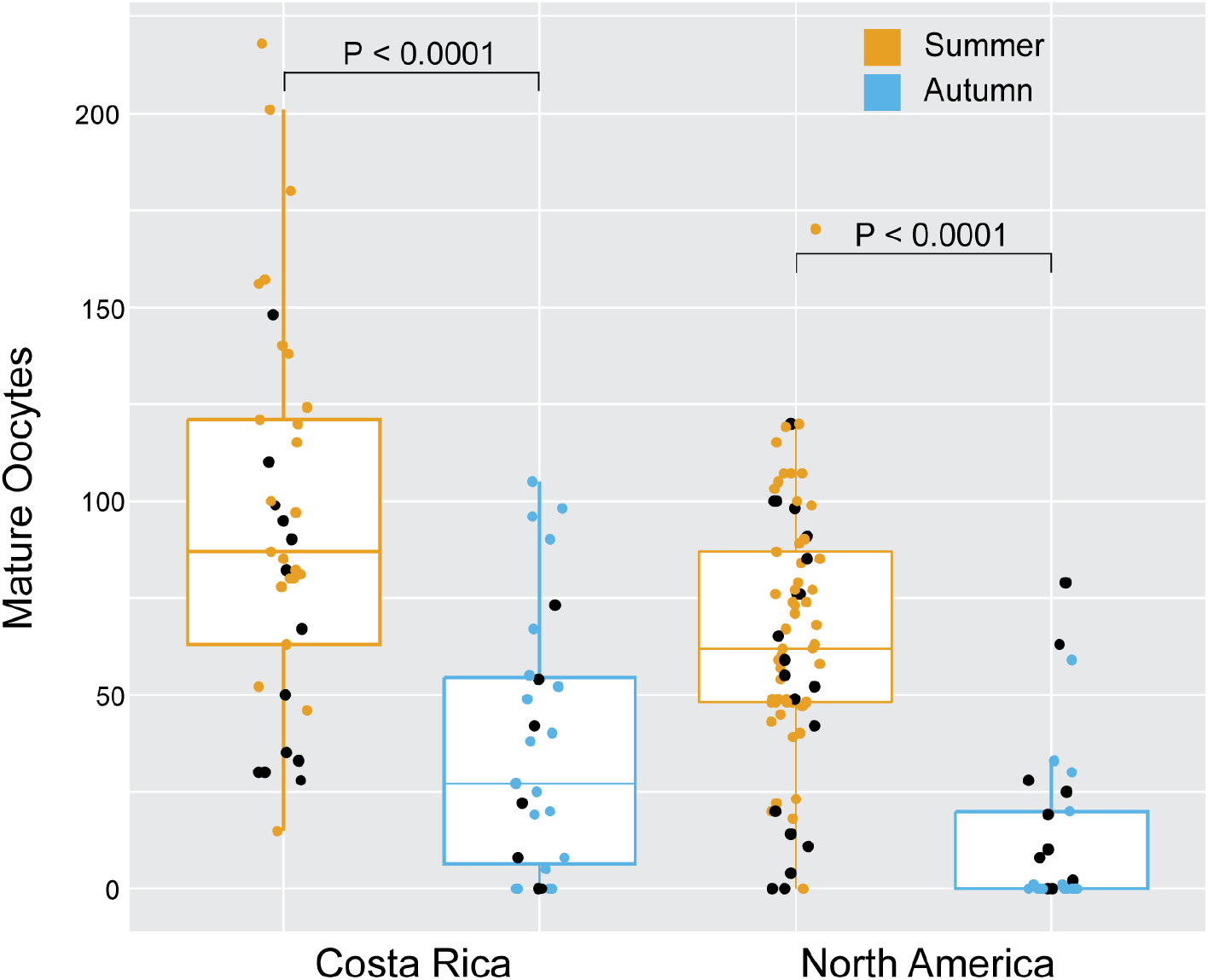
Boxplot of mature oocytes in female monarchs. Both CR and NA monarchs have decreased numbers of mature oocytes in response to autumn rearing, relative to summer rearing. Significant differences between rearing seasons are indicated on the plot. Black dots highlight individuals also assayed for metabolic rates.

### Components of body mass differ in seasonal plasticity between sexes and populations

Total body mass (the combined mass of the thorax and abdomen) did not differ significantly between populations, rearing seasons, or sexes (mean = 0.0845 grams, Kruskal-Wallis χ^2^= 11.4, df = 7, *P* = 0.12). However, abdomen mass was seasonally plastic in both NA and CR monarchs, and this plasticity differed between the sexes (Table 2). A model including season, sex, and their interaction explained ∼10% of variation in abdomen mass (R^2^ = 0.096; Table 2). Males reared in summer had lighter abdomens than females reared in summer (female mean = 0.0352g vs male mean = 0.043g, Kruskal-Wallis χ^2^= 16.8, df = 7, *P* = 0.019, Dunn test with Bonferroni correction, *P* = 0.0035), and males increased abdomen mass in response to autumn (autumn mean = .0454g vs summer mean = 0.0352g, Dunn test with Bonferroni correction, *P* = 0.0043).

**Table 2.**
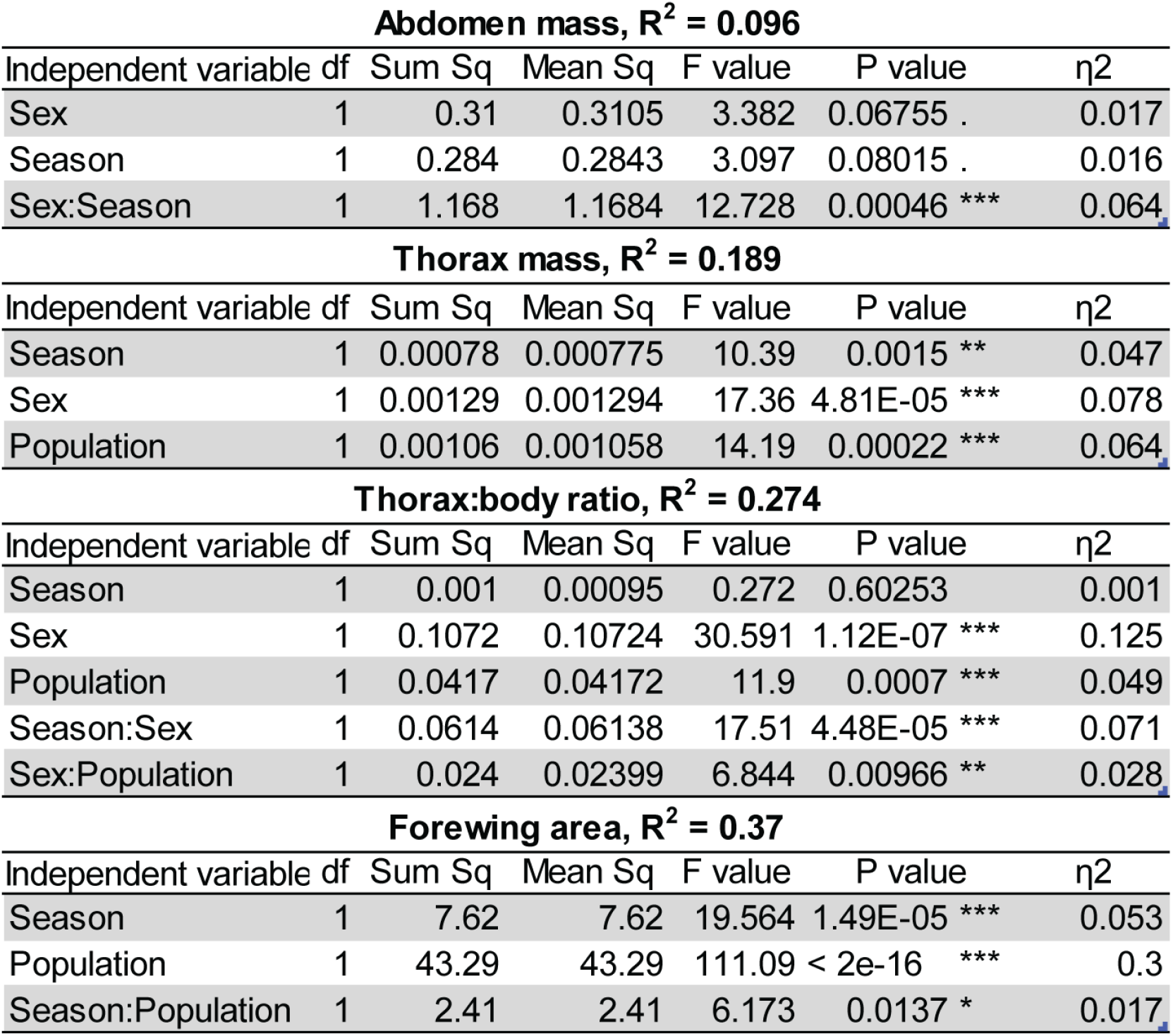
Summary of the best fit general linear model (glm) for abdomen mas, thorax mass, the ratio of thorax:body mass, and forewing area. Each model’s R^2^ is reported along with the significance and effect size of each independent variable in the model. η2 (Eta squared) is a measure of effect size that can be interpreted as the amount of variance accounted for by each variable in the best fit glm.

Thorax mass was seasonally plastic in both populations, with no evidence for a statistical interaction between season and population (Table 2). A model including rearing season, sex, and population explained 19% (R^2^ = 0.19) of the variation in thorax mass (Table 2). An individual’s thorax was likely to be heavier if population was NA, sex was male, and season of development was autumn (Supplementary Information, Figure S5). A post-hoc test found no significant differences between NA males and females or CR males and females reared in either season, though the difference between CR males and females was nearly significant in autumn (Kruskal-Wallis χ^2^= 38.67, df = 7, *P* < 0.0001, Dunn test with Bonferroni correction, NA: *P* = 0.88, *P* = 1.0, CR: *P* = 0.48, *P* = 0.08, summer and autumn respectively).

The ratio of thorax mass to total body mass was seasonally plastic in females, particularly in NA monarchs (Table 2). A model including population and sex, as well as interactions between population and sex and between sex and season explained 27% of the variation in the thorax:body mass ratio (R^2^ = 0.27). Sex had the largest effect with males having higher thorax:body mass ratios than females. NA monarchs had higher thorax:body mass ratios than did CR monarchs. Females increased the thorax:body mass ratio when reared in autumn relative to summer, and this effect of season was significant in NA but not in CR females (Figure 3, Kruskal-Wallis χ^2^= 52.97, df = 7, *P* < 0.0001, Dunn test with Bonferroni correction, CR: *P* = 1 and NA: *P* = 0.0246). Autumn-reared NA female thorax:body mass ratios were significantly greater than those of autumn-reared CR females (Figure 3, Dunn test with Bonferroni correction *P* = 0.0125). In summary, investment in thorax mass as a fraction of total body mass exhibits a sex-specific plasticity that was significant in NA monarchs, with NA females developing a more male-like pattern of investment in autumn versus summer.

**Figure 3.**
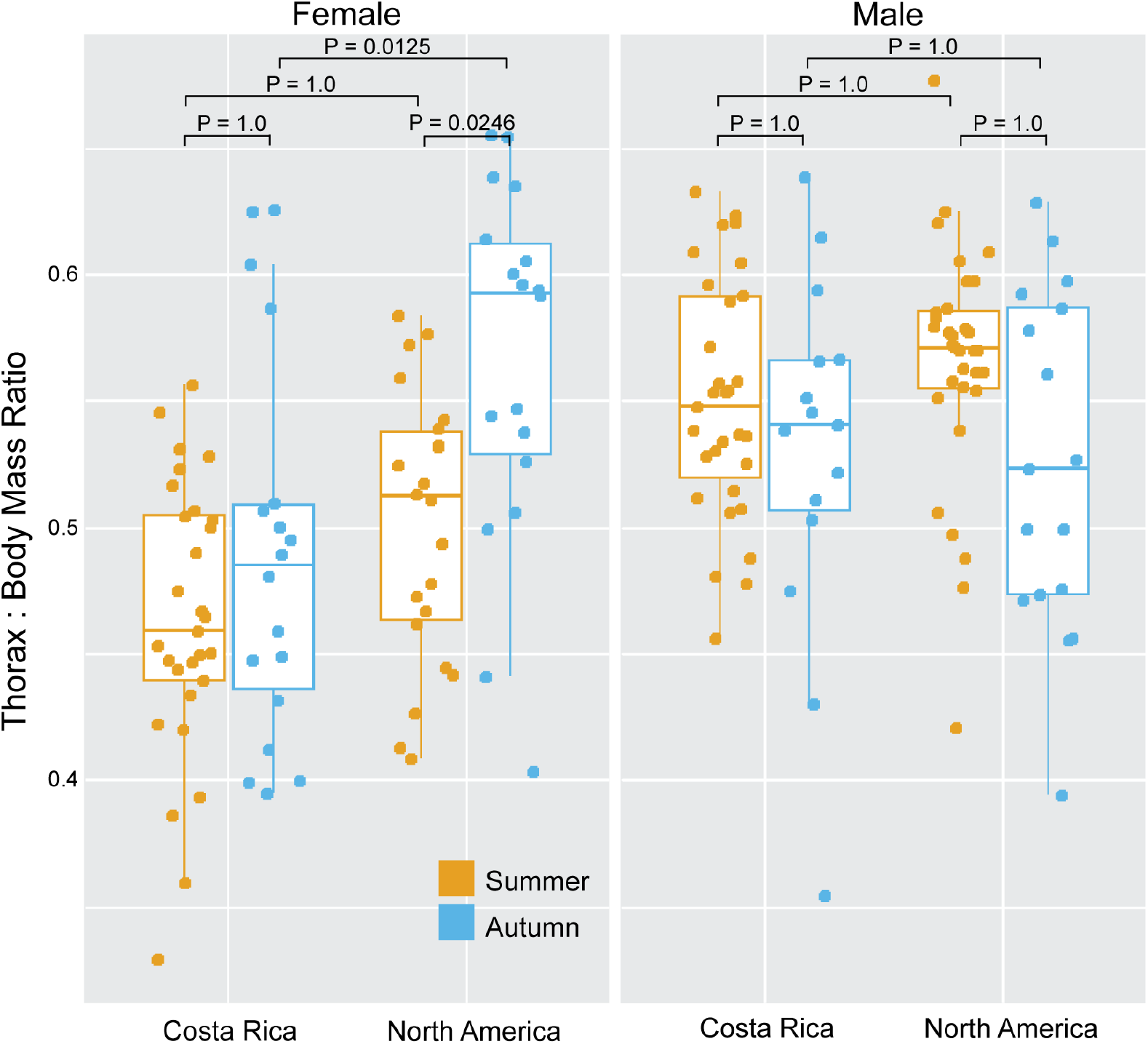
Boxplot of the ratio of thorax mass to total body (thorax + abdomen) mass. Scores above 0.5 indicate an individual has invested more of their total mass in the thorax than in abdomen. NA female monarchs increase investment in thorax tissue in autumn. P-values for differences between season, sex, and population are indicated on the plot.

### Only NA monarchs exhibit seasonal plasticity in wing size

Wing area was seasonally plastic in NA monarchs. Variation in wing area was best explained by a model that included the effects of rearing season and population, as well as their interaction (R^2^ = 0.37; Table 2). NA monarchs reared in autumn had on average 8% larger forewings than the NA summer-reared monarchs (Figure 4, summer mean =7.26 cm^2^ vs autumn mean = 7.87 cm^2^, Kruskal-Wallis χ^2^ = 90.68, df = 3, *P* < 0.0001, Dunn test with Bonferroni correction, *P* < 0.0001) and 16% larger forewings than the CR autumn-reared monarchs. CR monarch forewing area was not seasonally plastic (Figure 4, summer mean =6.56 cm^2^ vs autumn mean = 6.79 cm^2^, Dunn test with Bonferroni correction, *P* = 0.54).

**Figure 4.**
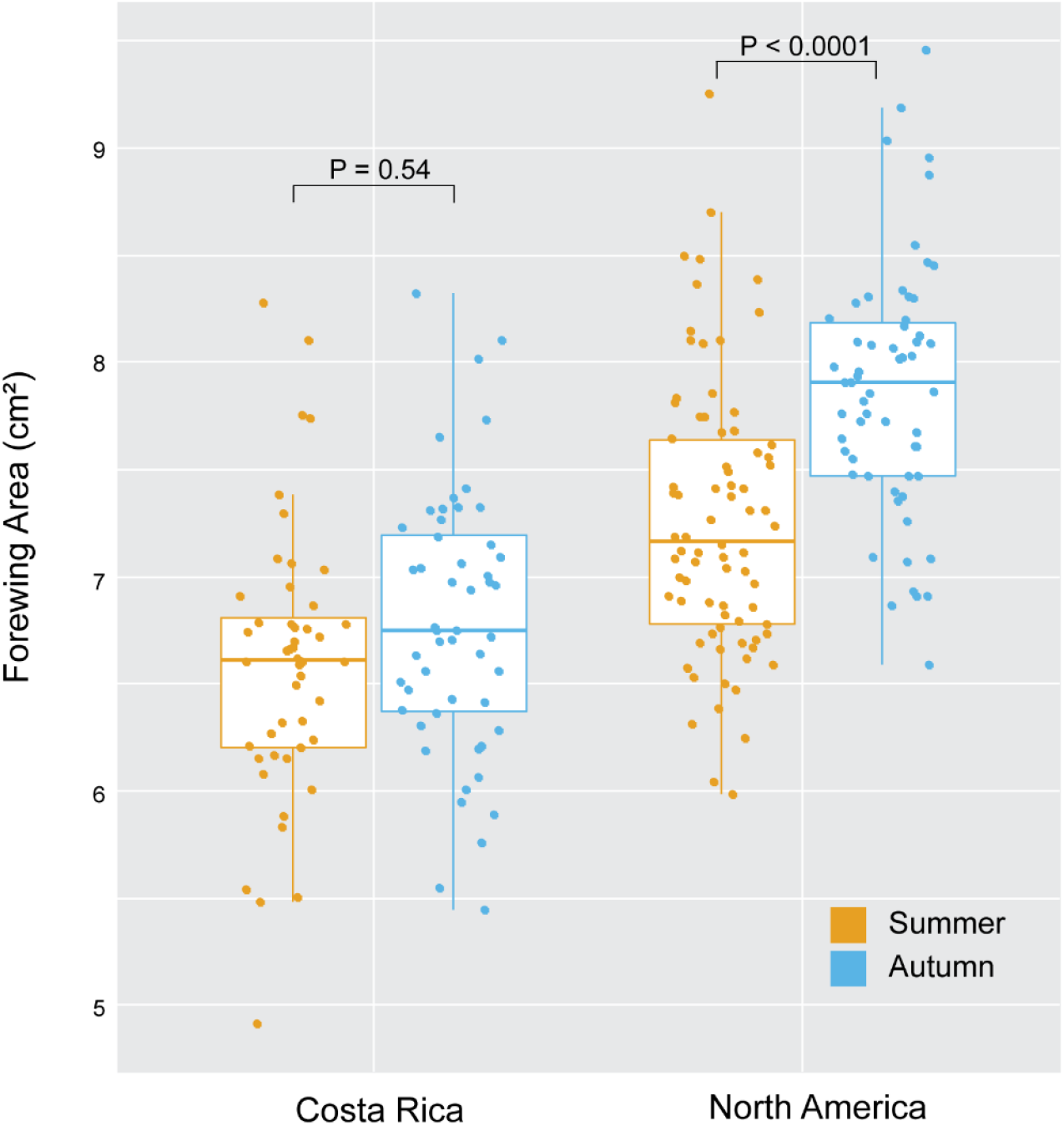
Boxplot of the forewing area measured in cm^2^. NA monarchs increase the size of their forewing in response to autumn while CR monarchs do not. P-values for differences between seasons in each population are indicated on the plot.

### Neither NA nor CR monarchs exhibit seasonal plasticity in wing shape

In contrast, measures of forewing shape did not differ between NA and CR monarchs and showed little to no within population plasticity. Variation in forewing aspect ratio was not explained by sex, season, population, or any of their interactions (Kruskal-Wallis χ^2^= 7.37, df = 7, *P* = 0.39). Circularity of the forewing, where a value of 1 is a perfect circle and decreasing scores indicate more polygonal (angular) forewings, was not seasonally plastic in either population, although NA monarchs trended towards more angular wings in autumn (Supplementary Information, Figure S8A, Kruskal-Wallis χ^2^= 8.72, df = 3, *P* = 0.03, Dunn test with Bonferroni correction, NA: *P* = 0.085 and CR: *P* = 0.85). Geometric morphometric analysis did not reveal any differences in mean shape between NA and CR forewings in either season, and neither population exhibited any seasonal plasticity in this measure of forewing shape (Supplementary Information, Figure S6 and S7). To quantify variability in forewing shape, we measured the distance of each individual forewing’s landmark to the respective consensus mean landmark and summed those distances. Total distance from the mean shape did not vary by population or season (Supplementary Information, Figure S8B, Kruskal-Wallis χ^2^= 3.58, df = 3, *P* = 0.31).

### Metabolic rates were seasonally plastic in CR but not NA monarchs

Resting MR of NA monarchs was not seasonally plastic (mass x rearing, *P* = 0.55; rearing, *P* = 0.37) (Figure 5B and Supplementary Information, Table S6). However, autumn-reared CR monarchs had significantly greater resting MR relative to summer-reared CR monarchs (mass x rearing, *P* = 0.86; rearing, *P* = 2.63E-08) (Figure 5A and Supplementary Information, Table S6). Variation in mass-corrected MR was explained by a significant interaction between rearing season and population (*P =* 0.004; Table 3), with only CR monarchs exhibiting seasonal plasticity (Figure 5C). Resting MR of summer-reared NA and CR monarchs were not significantly different, but resting MR of CR monarchs was significantly greater than that of NA monarchs reared in autumn (Supplementary Information, Table S7).

**Table 3.** Summary of general linear model used to test for effects of population and rearing season on mass-corrected MR.

| Trait | Independent Variable | df | Sum Sq | Mean Sq | F-value | P-value |
| --- | --- | --- | --- | --- | --- | --- |
| Mass-corrected resting MR | Population | 1 | 0.001 | 0.0014 | 0.03 | 0.8746 |
|  | Season | 1 | 0.879 | 0.8789 | 15.64 | 0.0001 *** |
|  | Population:Season | 1 | 0.490 | 0.4900 | 8.72 | 0.0036 ** |
| Mass-corrected flight MR | Population | 1 | 0.036 | 0.0360 | 0.70 | 0.404 |
|  | Season | 1 | 0.252 | 0.2515 | 4.89 | 0.0283 * |
|  | Population:Season | 1 | 0.023 | 0.0232 | 0.45 | 0.5026 |

**Figure 5.**
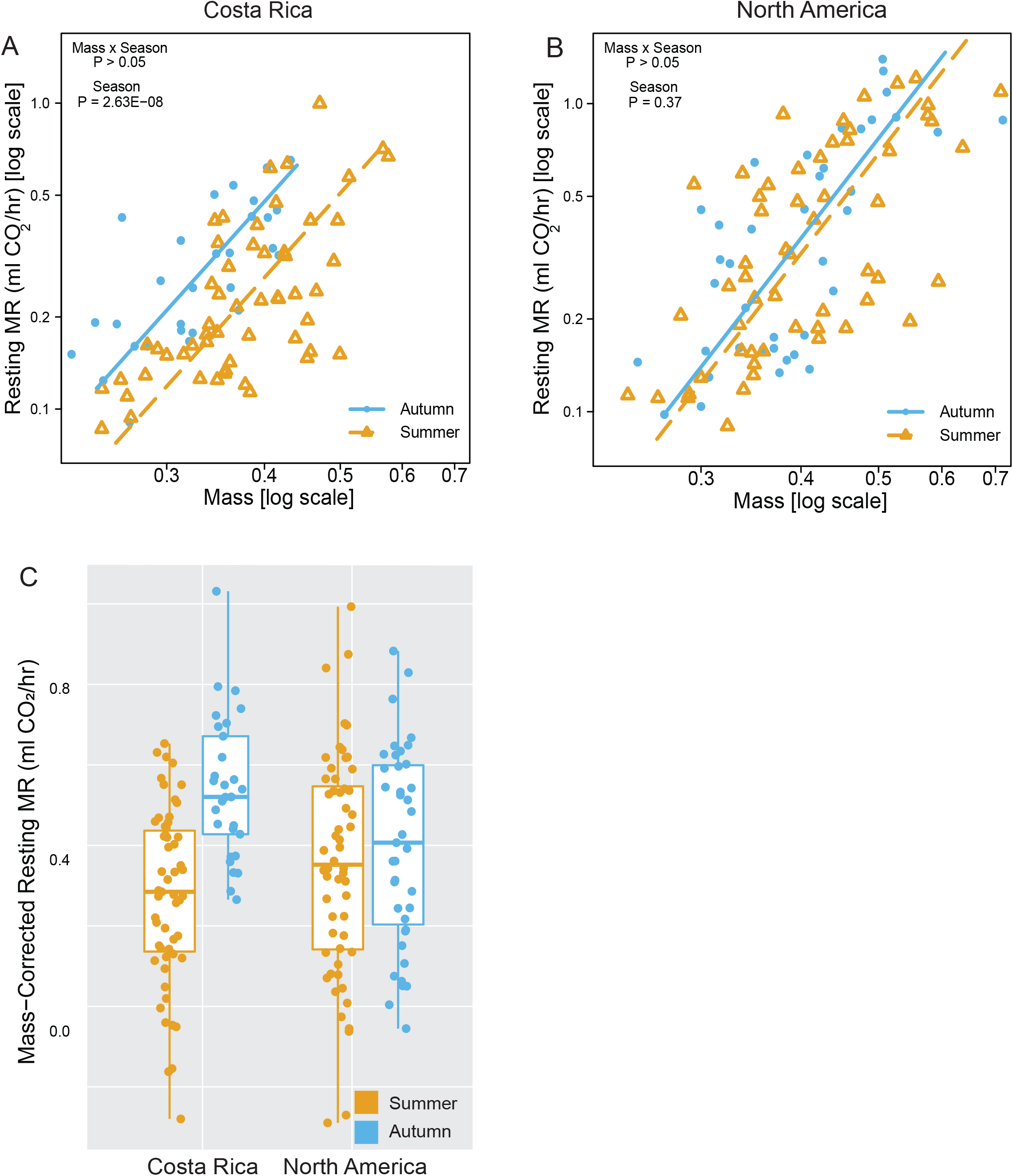
Effects of population and seasonal rearing conditions on resting metabolic rates (MR). A) Resting MR were significantly increased in CR monarchs reared in autumn relative to summer, B) while rearing season did not affect the resting MR of NA monarchs. C) There was a significant effect of the interaction between population and rearing season on mass-corrected resting MR (population x rearing, P = 0.004), with the maintenance of similar resting metabolic rates across seasonal rearing environments in NA but not in CR monarchs. Male and female data are plotted together, as the sexes did not differ in patterns of MR (Supplementary Information, Table S5).

Similar to patterns for resting MR, CR monarchs had elevated flight MR when reared in autumn relative to summer (mass x rearing, *P* = 0.19, rearing *P* = 0.04) (Figure 6A and Supplementary Information, Table S6), but maximal flight MR in NA monarchs was not seasonally plastic (mass x rearing, *P* = 0.97; rearing *P* = 0.19) (Figure 6B and Supplementary Information, Table S6). When we corrected flight MR for mass, there was a significant effect of season (*P =* 0.0283; Table 3) but no significant interaction between rearing season and population (Table 3). However, the magnitude of seasonal plasticity in mass-corrected flight MR appeared larger in CR relative to NA monarchs (Figure 6C). Flight MR of summer-reared NA and CR monarchs were not significantly different, but autumn-reared CR and NA monarchs differed in the scaling relationship with mass, with larger NA monarchs maintaining lower flight MR than CR monarchs (Supplementary Information, Table S7).

**Figure 6.**
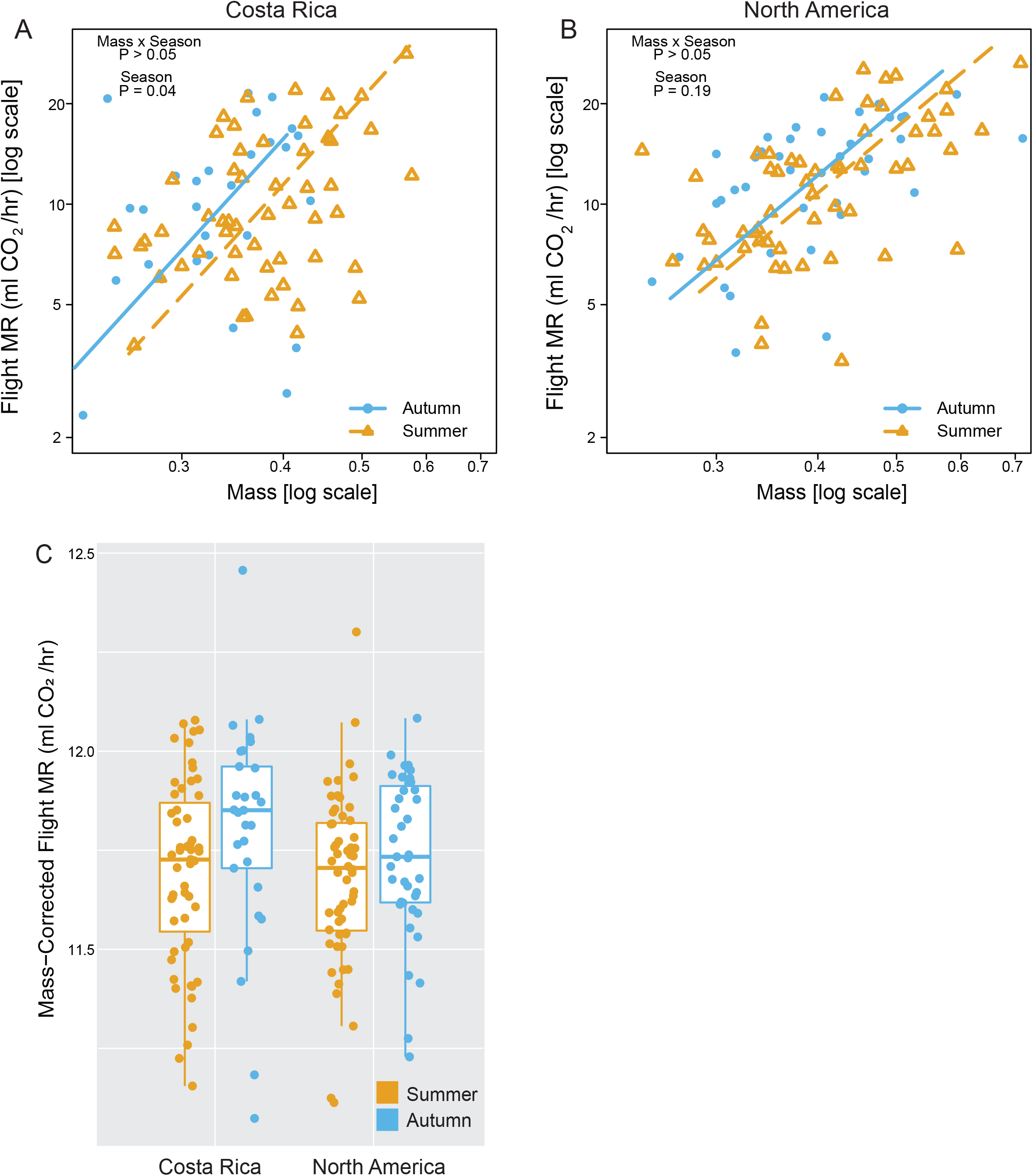
Effects of population and seasonal rearing conditions on flight metabolic rates (MR). A) CR monarchs had significantly greater flight MR when reared in autumn relative to summer. B) Flight MR were slightly elevated in, but not significantly different between autumn and summer-reared NA monarchs. C) Mass-corrected flight MR showed a similar pattern, with increased flight MR in autumn- relative to summer-reared monarchs (rearing, P = 0.03) and a larger magnitude of difference in CR monarchs. However, there was no statistically significant effect of the interaction (population x rearing, P = 0.50). Male and female data are plotted together, as the sexes did not differ in patterns of metabolic rate (Supplementary Information, Table S5).

## Discussion

We compared ancestral temperate (NA) and derived tropical (CR) monarch populations for the extent of seasonal plasticity in physiological and morphological traits suspected to be adaptive for monarch migration and overwintering. We predicted that plasticity would be lost in monarch populations that have dispersed into more stable, tropical habitats, such as Costa Rica. We found that the non-migratory CR descendants of the migratory NA population retain some, but not all ancestral seasonal trait plasticity. This suggests that seasonal plasticity in monarchs can be lost in a piecemeal fashion in the absence of selective pressures for its maintenance. The maintenance of metabolic rates in autumn compared to summer, plus the increase in wing size and thorax mass relative to total body mass in NA monarchs, suggest that these traits may be important for migration success and that the regulation of these traits may be critical to maintaining alternative summer and autumn phenotypes.

Mass differences in the abdomen and thorax are consistent with different selective pressures facing females versus males as well as NA versus CR populations. Autumn rearing induces an apparent shift in resources in females presumably from egg mass to flight muscle, consistent with the idea that successful autumn migration is critical for both sexes. While NA male and female monarchs were not significantly different in thorax mass in either season in our comparisons, a previous experiment that compared thorax mass in NA monarchs found significant differences between thorax mass in males and females (Davis and Holden 2015). Particularly in summer, we saw a similar trend towards larger male thoraxes, and sex was a significant predictor of thorax mass in our glm. Though not to the same degree as NA females, CR females also responded to autumn by increasing the thorax to body mass ratio though the difference comes from a decrease in abdomen mass rather than an increase in thorax mass in autumn. In summary, CR females retained seasonally plastic reproduction, but the seasonal shift in allocation to thorax mass may be eroding. Further investigation of plasticity in resource allocation into reproductive and flight muscle tissues are warranted, as well as investigation of whether other abiotic factors (e.g., drought or host-plant quality) may induce reproductive diapause and maintain plasticity for this trait in tropical monarch populations.

Forewing size was the most divergent morphological trait between NA and CR monarchs. Consistent with other work comparing migratory and resident monarch populations, we found CR monarchs had smaller wings than NA monarchs (Beall and Williams 1945; Dockx 2007; Altizer and Davis 2010; Li et al. 2016; Freedman et al. 2020). However, unlike previous work, our study explicitly compared monarchs reared in the NA monarch’s migratory range in summer and autumn. We found that forewing size was seasonally plastic in NA but not in CR monarchs. Previous measurements from a study of museum specimens collected in North America between 1878-2017 noted that autumn-collected individuals had larger wings than summer (Freedman and Dingle 2018). Our data suggest that this difference is at least partly explained by seasonal plasticity in wing size rather than differential mortality during migration (Flockhart et al. 2017; Davis et al. 2020). The smaller forewing size of CR and other resident monarch populations plus the CR monarchs’ lack of plasticity suggests that adaptation to the local environment post-dispersal may have selected for smaller wing size. Meanwhile, large wing size is likely under constant selection in migratory NA monarch populations during autumn, as large wing size is associated with longer flight in butterflies (Altizer and Davis 2010, Li et al. 2016, Flockhart et al. 2017). Thus, this might be an example where seasonal heterogeneity maintains plasticity in wing size in NA monarchs, with the summer-like small wing trait fixed in resident monarch populations that experience more summer-like conditions throughout the year. Investigating the flight and fitness consequences of these changes in wing morphology would be particularly useful for assessing whether this is an example of the loss of plasticity through adaptive assimilation.

The importance of wing shape to migration is less clear. Previous work found differences in shape between some resident and migratory monarchs (Dockx 2007; Altizer and Davis 2010; Satterfield and Davis 2014; Freedman et al. 2020), while other population comparisons did not find differences (Li et al. 2016; Freedman et al. 2020). Between our three measures of wing shape (geometric morphometrics, aspect ratio, and circularity), the only significant shape difference was in forewing circularity between autumn-reared NA monarchs and summer-reared CR monarchs, but the difference was small and the distributions were largely overlapping. We suggest that the difference seen in circularity when comparing wild-caught CR monarchs to NA monarchs (Altizer and Davis 2010) could be driven by developmental environment rather than population. However, we found no evidence of seasonal plasticity in wing shape in either population, consistent with findings from Flockhart et al. (2017) which found no relationship between wing roundness or aspect ratio and distance flown in NA migrators. However, others have noted differences in aspect ratio when comparing wild-caught to indoor-reared NA monarchs (Davis et al. 2020) and when comparing NA individuals caught earlier in the migration season to individuals caught later (Satterfield and Davis 2014).

In contrast to the prediction that NA monarchs relative to CR monarchs might exhibit greater plasticity in metabolic rates to support flight during migration, we observed that metabolic rates were affected by seasonal rearing only in CR monarchs. Autumn-reared CR monarchs had elevated resting and flight metabolic rates relative to summer-reared monarchs, while NA monarchs maintained similar and lower resting and flight metabolic rates across seasons. There are two, non-mutually exclusive, ways to interpret this pattern. First, the NA population may have seasonal plasticity in underlying physiology that maintains similar metabolic rates across seasonal environments, with the plastic mechanisms that maintain metabolic rate across the seasons lost in the CR population. Second, if the CR population has lost either the maternal provisioning or developmental mechanisms appropriate for the shorter photoperiod days of autumn, then the elevated metabolic rates in autumn-reared CR monarchs may be the consequence of coping with environmental stress during development. That stress, however, cannot be attributed to differences between reproductive output or host plant between the populations, as monarchs from both populations significantly decreased egg counts in response to autumn and consumed common milkweed in both summer and autumn in our common garden experiment. We note that while common milkweed differs from CR’s native tropical milkweed host (*Asclepias curassavica*), this did not result in differences in metabolic rate between the populations in the summer, suggesting that any effect of host plant on population differences in metabolic rate in our study must interact with the effect of seasonal rearing.

Our results were similar to previous studies of metabolic rates in NA and CR monarchs that were reared in summer (Zhan et al. 2014), although that study used somewhat different measures of metabolic rates and did detect differences in flight metabolic rate between migratory NA and resident Florida monarchs. Zhan et al. (2014) also found evidence for positive selection and divergent expression of collagen IV alpha-1 and alpha-2 in adult thoracic muscle tissue between migratory and non-migratory populations of monarchs. These proteins are essential for muscle morphogenesis and function (Schnorrer et al. 2010), and have been interpreted as evidence for the evolution of flight efficiency in migrating monarchs (Zhan et al. 2014). Flight is energetically demanding, and selection for long-distance migratory flight may favor more efficient flight relative to shorter duration flight (Rankin and Burchsted 1992). Our results lend support to this hypothesis, as we found that NA monarchs maintained similar resting and flight metabolic rates across seasons. We suggest that migration is supported not by increased metabolic output but likely through other seasonally plastic changes (e.g., in wing area, as we observed, and/or muscle structures) that enable more efficient flight. These results contrast with some other migratory and dispersing insects that have higher metabolic rates compared to their non-migratory and non-dispersing counterparts (Tanaka and Okuda 1996; Zera et al. 1997; Crnokrak and Roff 2002; Niitepõld et al. 2009). Of these examples, NA monarchs migrate the farthest and live the longest. Thus, the maintenance of low metabolic rates may enable monarchs to better survive the months-long overwintering period in Mexico where they consume very little food. Our observation that NA butterflies are able to maintain low flight MR unlike CR butterflies reared in autumn may also indicate that NA monarch physiology enables more efficient flight in the presence of accumulated lipid reserves during migration (Gibo and McCurdy 1993; Brower et al. 2006; Schroeder et al. 2020).

## Acknowledgements

We thank Michelle Noyes and Elena Sparrow for their help with rearing the monarchs in 2017. We also thank Trevor Price and Micah Freedman for constructive feedback on the manuscript as well as anonymous reviewers.

## Funding

This work was supported by the Graduate Research Fellowship Program, National Institute of Health Genetics and Regulation Training Grant (T32 GM07197), US Fish and Wildlife Service (Award F17AC01222), National Science Foundation Grants (IOS-1452648 and IOS-1736249), and National Institute of Health Grant (GM131828).

## Conflict of Interest Statement

The authors declare no conflicts of interest.

## Author Contributions

All authors participated in conceiving the ideas and designing methodology; AT-T, WL and CRJ collected the data; AT-T, CRJ and KLM analysed the data; AT-T and CRJ led the writing of the manuscript. All authors contributed critically to the drafts and gave final approval for publication.

## Data Availability

All data are included in supplemental file Data.xlsx.

## Notes

### Competing Interest Statement

The authors have declared no competing interest.

