## Supplementary Information for "Seasonal plasticity in morphology and metabolism differs between migratory North American and resident Costa Rican monarch butterflies"

*Supplemental Figures*

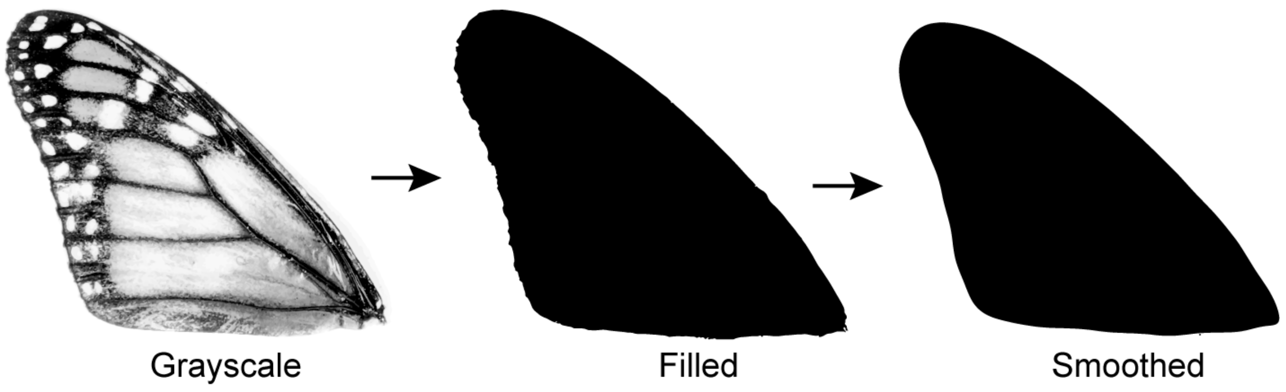


**Figure S1**. Example of a monarch forewing being processed in ImageJ. The photo is converted to an 8-bit black/white image. The non-black pixels within the forewing are filled with black. Once filled, the edges of the wing are smoothed using the ‘Shape smoothing’ plugin.

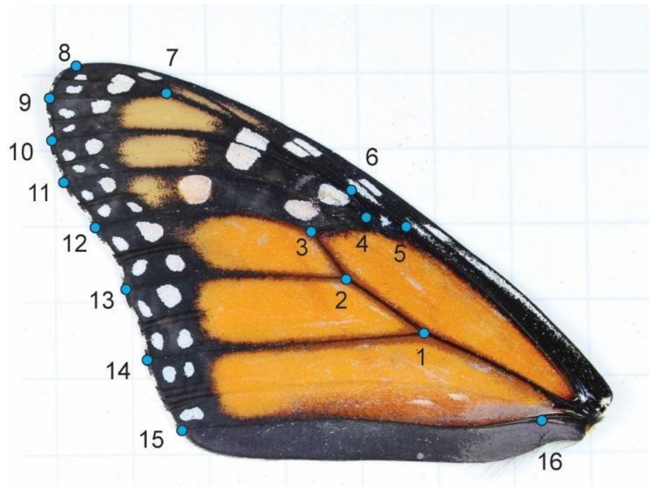


**Figure S2**. Blue dots indicate the positions of the 16 landmarks on the monarch butterfly forewing.

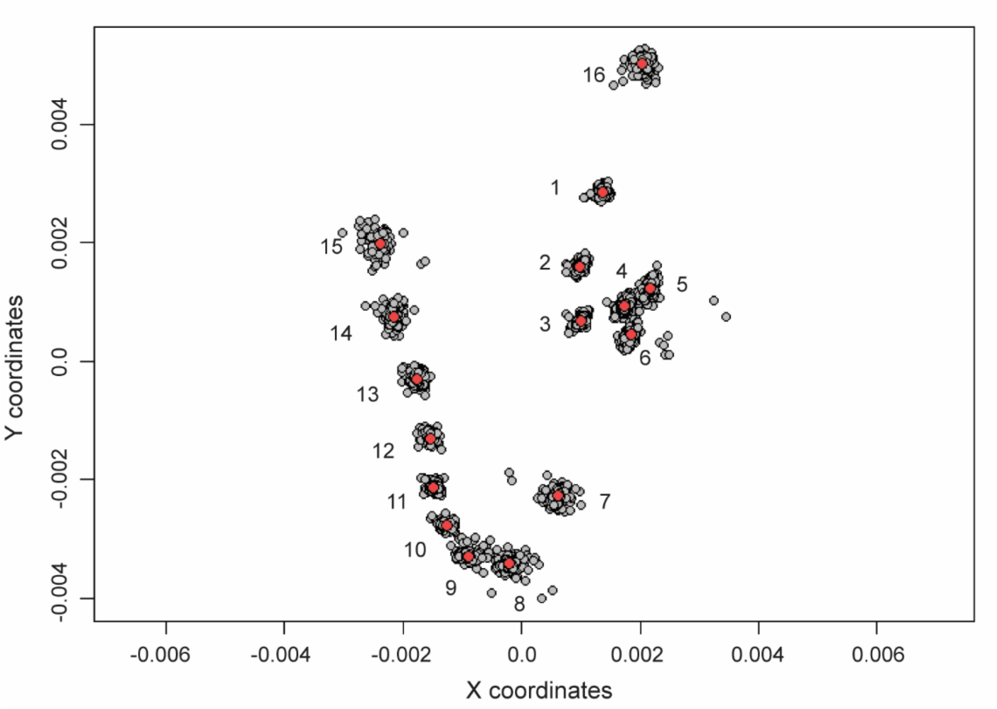


**Figure S3**. Plot of all specimens' landmarks after general Procrustes alignment in grey. Red dots are the consensus mean coordinates for each landmark. Landmarks numbered 1 through 16 correspond to vein intersections and margins in figure S2.

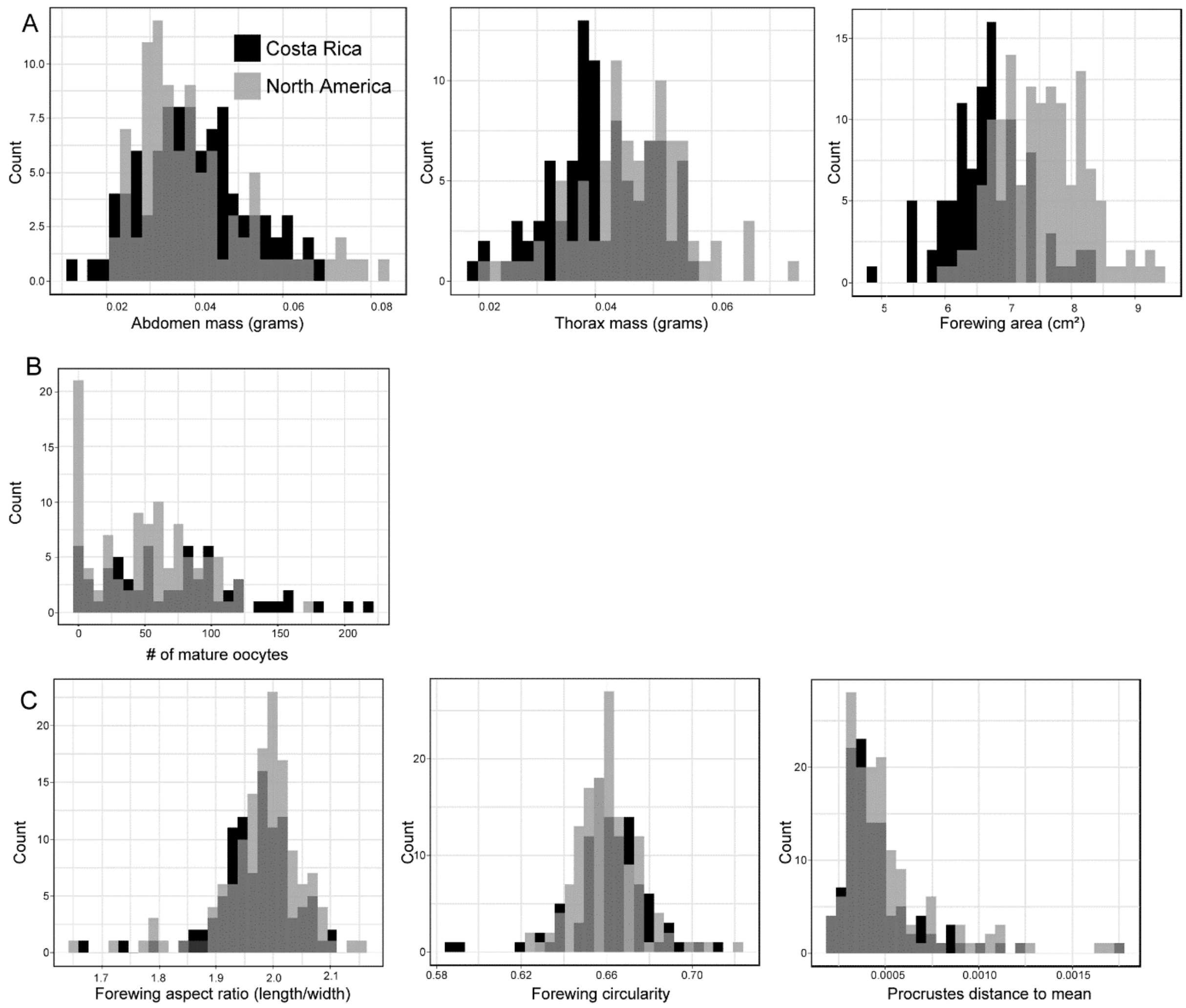


**Figure S4**. Distributions of morphological phenotypes in CR and NA populations. A) Abdomen mass, thorax mass, and forewing area are all normally distributed while B) the number of mature oocytes follows a negative binomial distribution. C) Forewing shape scores including aspect ratio, circularity, and Procrustes distance from the mean shape are not normally distributed.

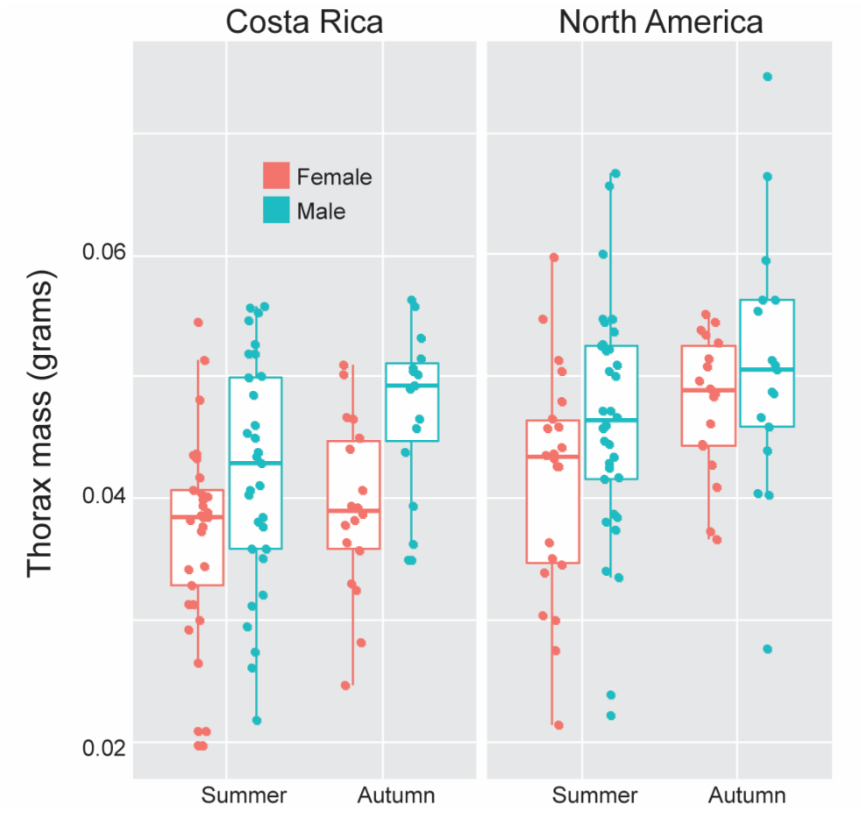


**Figure S5**. Boxplot of thorax mass measured in grams.

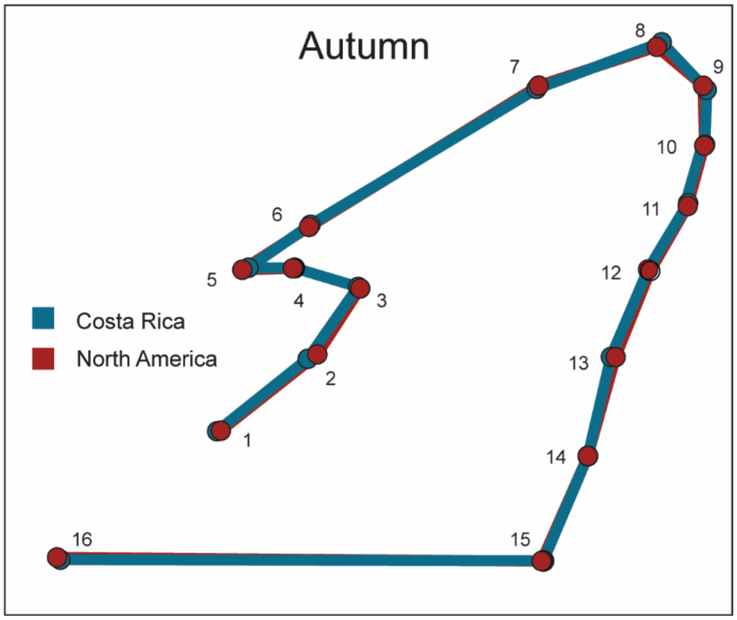


**Figure S6**. Comparison of mean forewing shape of Costa Rican (blue) and North American (red) monarchs reared outdoors in autumn. Each dot is the consensus mean coordinate for landmarks 1-16. Straight lines are drawn between landmarks to outline the forewing.

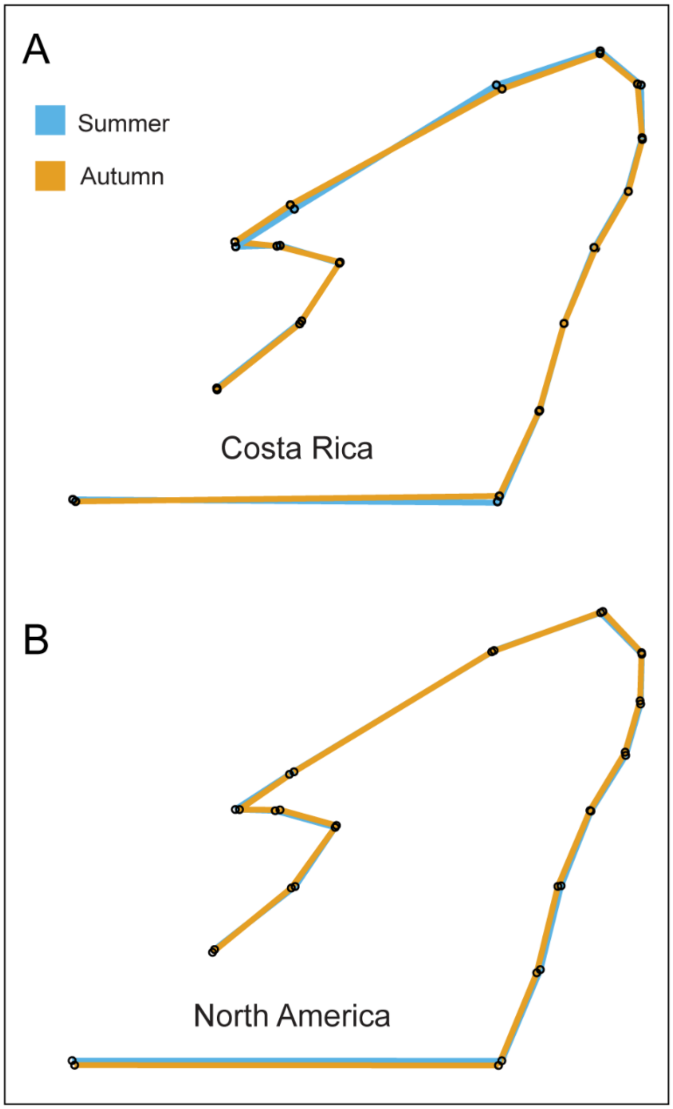


**Figure S7**. Comparison of mean forewing shapes of Costa Rican and North American monarchs reared outdoors in summer and autumn. Each dot is the consensus mean coordinate for landmarks 1-16. Straight lines are drawn between landmarks to outline the forewing. A) Mean shape of the Costa Rican autumn (yellow) forewing plotted on top of the mean shape of Costa Rican summer (blue) forewing. B) Mean shape of the North American autumn forewing (yellow) plotted on top of the mean North American summer (blue) forewing.

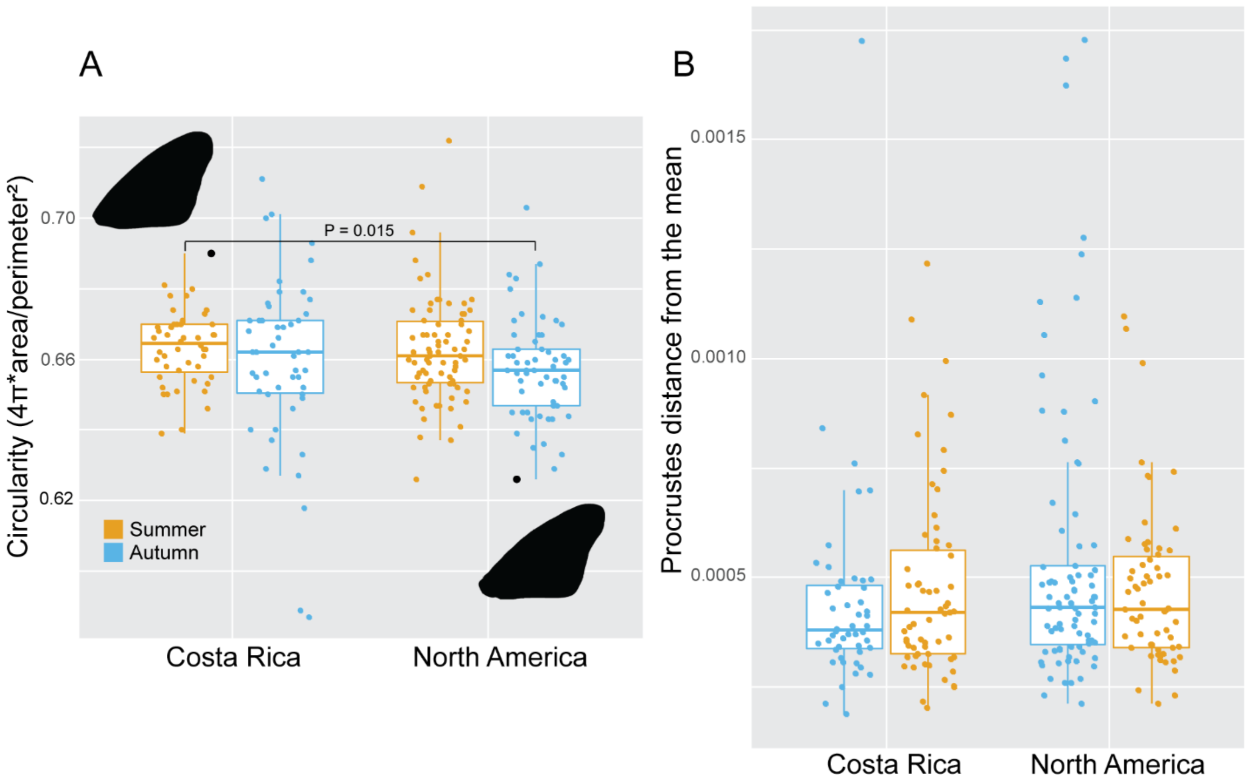


**Figure S8.** A) Boxplot of forewing circularity scores. Lower circularity scores indicate an individual with a more elongated wing. CR monarchs reared in summer had higher circularity scores than NA monarchs reared in autumn. The p-value of the only significant difference between groups is indicated on the plot. Examples of both a less elongated CR monarch wing and a more elongated NA monarch wing are highlighted in black. B) A boxplot of Procrustes distances from the mean consensus forewing shape. Each dot represents the cumulative distance of 16 coordinates (landmarks) from their respective mean shape coordinate. There were no significant differences between groups.

*Supplemental Tables*

**Table S1.** Combined sample sizes for metabolic rate measurements of monarchs reared in 2016 and 2017.

| | Sample sizes 2016 + 2017 (N = 179) | | | |
|---|---|---|---|---|
| | Summer (N =111) | | Autumn (N = 68) | |
| | Females | Males | Females | Males |
| North American | 29 | 27 | 16 | 23 |
| Costa Rican | 32 | 23 | 14 | 15 |

**Table S2.** Candidate general linear models for mass measurements. Models were fit with Gaussian distribution and ranked by BIC score and model weight. Best fit model for each dependent variable is highlighted in red.

| Abdomen mass candidate models | BIC | Weight |
|---|---|---|
| Abdomen mass ~ 1 + Season + Sex+ SexSeason | 104.784 | 0.6458 |
| Abdomen mass ~ 1 | 107.794 | 0.1434 |
| Abdomen mass ~ 1 + Season | 109.598 | 0.0582 |
| Abdomen mass ~ 1 + Sex | 109.858 | 0.0511 |
| Abdomen mass ~ 1 + Season + Sex+ Population + Sex Season | 109.994 | 0.0477 |
| Abdomen mass ~ 1 + Season + Sex | 112.140 | 0.0163 |
| Abdomen mass ~ 1 + Season + Sex+ Population + Sex Season + Population:Sex | 112.875 | 0.0113 |
| Abdomen mass ~ 1 + Population | 113.001 | 0.0106 |
| Abdomen mass ~ 1 + Season + Population | 114.792 | 0.0043 |
| Abdomen mass ~ 1 + Season + Sex+ Population + Sex Season + Population:Season | 115.030 | 0.0038 |
| Abdomen mass ~ 1 + Sex+ Population | 115.072 | 0.0038 |
| Abdomen mass ~ 1 + Season + Sex+ Population | 117.354 | 0.0012 |
| Abdomen mass ~ 1 + Sex+ Population + Population:Sex | 117.901 | 0.0009 |
| Abdomen mass ~ 1 + Season + Sex+ Population + Sex Season + Population:Season + Population:Sex | 118.007 | 0.0009 |
| Abdomen mass ~ 1 + Season + Sex+ Population + Population:Sex | 119.971 | 0.0003 |
| Abdomen mass ~ 1 + Season + Population + Population:Season | 119.976 | 0.0003 |
| Abdomen mass ~ 1 + Season + Sex+ Population + Population:Season | 122.519 | 0.0001 |
| Abdomen mass ~ 1 + Season + Sex+ Population + Population:Season + Population:Sex | 125.180 | 0.0000 |

| Thorax mass candidate models | BIC | Weight |
|---|---|---|
| Thorax mass ~ 1 + Season + Sex+ Population | -1204.551 | 0.7300 |
| Thorax mass ~ 1 + Season + Sex+ Population + Population:Sex | -1199.945 | 0.0730 |
| Thorax mass ~ 1 + Season + Sex+ Population + Population:Season | -1199.904 | 0.0715 |
| Thorax mass ~ 1 + Season + Sex+ Population + Sex Season | -1199.461 | 0.0573 |
| Thorax mass ~ 1 + Sex+ Population | -1198.101 | 0.0290 |
| Thorax mass ~ 1 + Season + Sex | -1195.801 | 0.0092 |
| Thorax mass ~ 1 + Season + Sex+ Population + Population:Season + Population:Sex | -1195.203 | 0.0068 |
| Thorax mass ~ 1 + Season + Sex+ Population + Sex Season + Population:Sex | -1194.878 | 0.0058 |
| Thorax mass ~ 1 + Season + Population | -1194.865 | 0.0058 |
| Thorax mass ~ 1 + Season + Sex+ Population + Sex Season + Population:Season | -1194.788 | 0.0055 |
| Thorax mass ~ 1 + Sex+ Population + Population:Sex | -1193.626 | 0.0031 |
| Thorax mass ~ 1 + Population | -1191.250 | 0.0009 |
| Thorax mass ~ 1 + Season + Sex+ Sex Season | -1190.640 | 0.0007 |
| Thorax mass ~ 1 + Season + Sex+ Population + Sex Season + Population:Season + Population:Sex | -1190.107 | 0.0005 |
| Thorax mass ~ 1 + Season + Population + Population:Season | -1190.027 | 0.0005 |
| Thorax mass ~ 1 + Sex | -1189.298 | 0.0004 |
| Thorax mass ~ 1 + Season | -1185.265 | 0.0000 |
| Thorax mass ~ 1 | -1181.654 | 0.0000 |

| Thorax:Body mass ratio candidate models | BIC | Weight |
|---|---|---|
| Thorax Body ~ 1 + Season + Sex+ Population + Sex Season + Population:Sex | -487.649 | 0.6113 |
| Thorax Body ~ 1 + Season + Sex+ Population + Sex Season | -485.922 | 0.2578 |
| Thorax Body ~ 1 + Season + Sex+ Population + Sex Season + Population:Season + Population:Sex | -482.772 | 0.0534 |
| Thorax Body ~ 1 + Season + Sex+ Population + Sex Season + Population:Season | -481.361 | 0.0263 |
| Thorax Body ~ 1 + Sex+ Population + Population:Sex | -481.047 | 0.0225 |
| Thorax Body ~ 1 + Season + Sex+ Sex Season | -480.441 | 0.0166 |
| Thorax Body ~ 1 + Sex+ Population | -479.008 | 0.0081 |
| Thorax Body ~ 1 + Season + Sex+ Population + Population:Sex | -476.415 | 0.0022 |
| Thorax Body ~ 1 + Season + Sex+ Population | -474.484 | 0.0008 |
| Thorax Body ~ 1 + Sex | -473.543 | 0.0005 |
| Thorax Body ~ 1 + Season + Sex+ Population + Population:Season + Population:Sex | -471.307 | 0.0002 |
| Thorax Body ~ 1 + Season + Sex+ Population + Population:Season | -469.580 | 0.0001 |
| Thorax Body ~ 1 + Season + Sex | -469.184 | 0.0001 |
| Thorax Body ~ 1 + Population | -461.033 | 0.0000 |
| Thorax Body ~ 1 + Season + Population | -455.948 | 0.0000 |
| Thorax Body ~ 1 | -454.852 | 0.0000 |
| Thorax Body ~ 1 + Season + Population + Population:Season | -450.882 | 0.0000 |
| Thorax Body ~ 1 + Season | -449.842 | 0.0000 |

**Table S3.** Candidate general linear model for forewing area. Models fit with Gaussian distribution and ranked by BIC score and model weight. Best fit model is highlighted in red.

| Forewing area candidate models | BIC | Weight |
|---|---|---|
| Forewing Area ~ 1 + Season + Population + Population:Season | 472.549 | 0.3379 |
| Forewing Area ~ 1 + Season + Population | 473.279 | 0.2346 |
| Forewing Area ~ 1 + Season + Sex + Population | 473.646 | 0.1952 |
| Forewing Area ~ 1 + Season + Sex + Population + Population:Season | 473.800 | 0.1808 |
| Forewing Area ~ 1 + Season + Sex + Population + Population:Sex | 478.988 | 0.0135 |
| Forewing Area ~ 1 + Season + Sex + Population + Sex:Season | 479.092 | 0.0128 |
| Forewing Area ~ 1 + Season + Sex + Population + Population:Season + Population:Sex | 479.262 | 0.0118 |
| Forewing Area ~ 1 + Season + Sex + Population + Sex:Season + Population:Season | 479.267 | 0.0117 |
| Forewing Area ~ 1 + Season + Sex + Population + Sex:Season + Population:Sex | 484.423 | 0.0009 |
| Forewing Area ~ 1 + Season + Sex + Population + Sex:Season + Population:Season + Population:Sex | 484.729 | 0.0008 |
| Forewing Area ~ 1 + Population | 494.484 | 0.0000 |
| Forewing Area ~ 1 + Sex + Population | 497.180 | 0.0000 |
| Forewing Area ~ 1 + Sex + Population + Population:Sex | 502.619 | 0.0000 |
| Forewing Area ~ 1 + Season | 558.223 | 0.0000 |
| Forewing Area ~ 1 + Season + Sex | 562.388 | 0.0000 |
| Forewing Area ~ 1 | 565.636 | 0.0000 |
| Forewing Area ~ 1 + Season + Sex + Sex:Season | 567.363 | 0.0000 |
| Forewing Area ~ 1 + Sex | 570.446 | 0.0000 |

**Table S4.** Candidate generalized linear model for mature oocyte counts. Models were fit with negative binomial distribution and ranked by BIC score and model weight. The best fit model is highlighted in red.

| Mature oocyte count candidate models | BIC | Weight |
|---|---|---|
| Oocytes ~ 1 + Population + Season | 1650.049 | 0.3694 |
| Oocytes ~ 1 + Population + Season + Season:Population | 1651.738 | 0.1588 |
| Oocytes ~ 1 + Population + Season + Year + Year:Population | 1651.963 | 0.1419 |
| Oocytes ~ 1 + Population + Season + Year + Year:Population + Year:Season | 1652.033 | 0.1370 |
| Oocytes ~ 1 + Population + Season + Year | 1652.428 | 0.1124 |
| Oocytes ~ 1 + Population + Season + Year + Season:Population | 1655.801 | 0.0208 |
| Oocytes ~ 1 + Population + Season + Year + Year:Season | 1655.958 | 0.0193 |
| Oocytes ~ 1 + Population + Season + Year + Season:Population + Year:Population | 1656.758 | 0.0129 |
| Oocytes ~ 1 + Season | 1657.136 | 0.0107 |
| Oocytes ~ 1 + Population + Season + Year + Season:Population + Year:Population + Year:Season | 1657.145 | 0.0106 |
| Oocytes ~ 1 + Population + Season + Year + Season:Population + Year:Season | 1658.811 | 0.0046 |
| Oocytes ~ 1 + Season + Year | 1661.212 | 0.0014 |
| Oocytes ~ 1 + Season + Year + Year:Season | 1666.230 | 0.0001 |
| Oocytes ~ 1 + Population + Year + Year:Population | 7214.004 | 0.0000 |
| Oocytes ~ 1 + Population + Year | 7230.434 | 0.0000 |
| Oocytes ~ 1 + Population | 7236.518 | 0.0000 |
| Oocytes ~ 1 | 7500.162 | 0.0000 |
| Oocytes ~ 1 + Year | 7504.577 | 0.0000 |

**Table S5.** Summary of a standardized major axis regression (sma) used to fit the metabolic scaling relations between ln (VCO2) and ln (mass) to test for effects of sex on metabolic rate (MR).

| Season, Trait | Population | Sex | $H_0$: equal slopes Slope (95% CI)[1] | $H_0$: no elevation difference Y-intercept (95% CI)[2] |
|---|---|---|---|---|
| Summer, Resting MR | NA | | LR = 0.006, df = 1, *P* = 0.93 | Wald = 2.82, df = 1, *P* = 0.09 |
| | | | Common Slope: 3.17 (2.56, 3.92) | |
| | | Male | | 0.75 (0.45, 1.06) |
| | | Female | | 0.79 (0.50, 1.08) |
| | Costa Rica | | LR = 0.64, df = 1, *P* = 0.42 | Wald = 0.14, df = 1, *P* = 0.70 |
| | | | Common Slope: 2.81 (2.30, 3.46) | |
| | | Male | | 0.53 (0.28, 1.79) |
| | | Female | | 0.55 (0.29, 0.81) |
| Autumn, Resting MR | NA | | LR = 1.11 df = 1, *P* = 0.29 | Wald = 0.11, df = 1, P = 0.73 |
| | | | Common Slope: 3.59 (2.80, 4.52) | |
| | | Male | | 0.98 (0.60, 1.36) |
| | | Female | | 1.01 (0.64, 1.37) |
| | Costa Rica | | LR = 0.13, df = 1, *P* = 0.71 | Wald = 0.23, df = 1, *P* = 0.63 |
| | | | Common Slope: 2.79 (2.12, 3.67) | |
| | | Male | | 0.80 (0.40, 1.20) |
| | | Female | | 0.77(0.36, 1.17) |
| Summer, Max flight MR | N. America | | LR = 0.17, df = 1, *P* = 0.67 | Wald = 3.79, df = 1, *P* = 0.09 |
| | | | Common Slope: 2.06 (1.64, 2.58) | |
| | | Male | | 1.90 (1.69, 2.12) |
| | | Female | | 1.80 (1.61, 1.99) |
| | Costa Rica | | LR = 0.64, df = 1, *P* = 0.42 | Wald = 2.73, df = 1, *P* = 0.85 |
| | | | Common Slope: 2.31 (1.78, 2.99) | |
| | | Male | | 2.03 (1.74, 2.32) |
| | | Female | | 1.92 (1.65, 2.19) |
| Autumn, Max flight MR | N. America | | LR = 2.743, df = 1, *P* = 0.09) | Wald = 0.07, df = 1, *P* = 0.78 |
| | | | Common Slope: 1.90 (1.42, 2.58) | |
| | | Male | | 1.84 (1.59, 2.09) |
| | | Female | | 1.83 (1.59, 2.06) |
| | Costa Rica | | LR = 0.03, df = 1, *P* = 0.84 | Wald = 0.2.28, df = 1, *P* = 0.13 |
| | | | Common Slope: 3.00 (2.00, 4.47) | |
| | | Male | | 2.50 (1.87, 3.13) |
| | | Female | | 2.32 (1.67, 2.98) |

**Table S6.** Summary of a standardized major axis regression (sma) used to fit the metabolic scaling relations between ln (VCO2) and ln (mass), and to test for effects of rearing season on metabolic rate (MR) within populations.

| **Population, Trait** | **Season** | **$H_0$: equal slopes Slope (95% CI)[1]** | **$H_0$: no elevation difference Y-intercept[2]** |
|---|---|---|---|
| Costa Rica, resting MR | | LR = 0.03, df = 1, *P* = 0.86 Common Slope: 2.84 (2.41, 3.34) | Wald = 30.96, df = 1, *P* = 2.63E-08 |
| | Autumn | | 0.81*(0.41, 1.16) |
| | Summer | | 0.56*(0.31, 0.83) |
| N. America, resting MR | | LR = 0.35, df = 1, *P* = 0.55 Common Slope: 3.34 (2.86, 3.89) | Wald = 0.78, df = 1, *P* = 0.37 |
| | Autumn | | 0.89 (0.66, 1.12) |
| | Summer | | 0.84 (0.62, 1.06) |
| Costa Rica, max flight MR | | LR = 1.79, df = 1, *P* = 0.19 Common Slope: 2.20 (1.85, 2.61) | Wald = 4.13, df = 1, *P* = 0.04 |
| | Autumn | | 2.25*(1.95, 2.55) |
| | Summer | | 2.12*(1.87, 2.37) |
| N. America, max flight MR | | LR = 0.00, df = 1, *P* = 0.97 Common Slope: 2.03 (1.70, 2.42) | Wald = 1.69, df = 1, *P* = 0.19 |
| | Autumn | | 1.89 (1.73, 2.05) |
| | Summer | | 1.84 (1.69, 1.99) |

[1]Test of common slope from a Type II regression model
[2]Significant differences in y-intercept within a common slope are evidence of differences in MR across the range of masses measured

**Table S7.** Summary of a standardized major axis regression (sma) used to fit the metabolic scaling relations between ln (VCO2) and ln (mass), and to test for effects of population on metabolic rate (MR) within seasons.

| **Season, Trait** | **Population** | **$H_0$: equal slopes Slope (95% CI)[1]** | **$H_0$: no elevation difference Y-intercept[2]** |
|---|---|---|---|
| Summer, resting MR | | LR = 0.53, df = 1, $P$ = 0.46<br>Common slope: 3.03 (2.62, 3.51) | Wald = 2.82, df = 1, $P$ = 0.09 |
| | N. America | | 0.72 (0.53, 0.91) |
| | Costa Rica | | 0.64 (0.44,0.84) |
| Autumn, resting MR | | LR = 1.68, df = 1, $P$ = 0.19<br>Common slope: 3.18 (2.67, 3.81) | Wald = 6.597, df = 1, $P$ = 0.01) |
| | N. America | | 0.82*(0.62, 1.31) |
| | Costa Rica | | 0.97*(0.41, 1.16) |
| Summer, max flight MR | | LR = 1.01, df = 1, $P$ = 0.31<br>Common slope: 2.20 (1.85, 2.61) | Wald = 0.10, df = 1, $P$ = 0.74 |
| | N. America | | 1.91 (1.75, 2.06) |
| | Costa Rica | | 1.92 (1.75, 2.09) |
| Autumn, max flight MR | | LR = 4.09, df = 1, $P$ = 0.04 | |
| | N. America | 2.02 (1.53, 2.67) | |
| | Costa Rica | 3.28 (2.24, 4.79) | |

[1]Test of common slope from a Type II regression model
[2]Significant differences in y-intercept within a common slope are evidence of differences in MR across the range of masses measured
